# Tuning evolutionary information in MSAs shapes protein generation and folding

**DOI:** 10.64898/2026.11.23.768656

**Authors:** Graham Richardson, Lionel Colliandre

## Abstract

Protein multiple sequence alignments are widely regarded as fixed artifacts of homologue retrieval. An alignment’s row composition shapes the evolutionary context that is supplied to a protein model. In this study, we tested whether tuning row composition would steer protein generation and structure prediction outcomes. We tuned lower-tail alignment information by controlling row selection within a reference protein-projected coordinate frame. We paired each information-targeted alignment with a matched-depth random-row control alignment. Across a cohort of 45 diverse proteins, row coherence among the retained homologues is predictive of structural confidence readouts and generated sequence novelty. The strongest row-coherence states occur near an empirical lower-tail information reference point of 1 bit/site. Below this reference point, generated de novo sequences demonstrate more eager exploration, higher decoder uncertainty, and reduced structural confidence at mutated positions. Across an independent structure-prediction benchmark, controlling the fraction of low-information positions demonstrated that intermediate fractions of low-information positions were associated with improved structure prediction within a confidence-gated subset. These findings evince that evolutionary information in protein alignments can be tuned to shape protein generation and structure prediction results.

## Introduction

The naturally occurring variation across protein evolution operates within an equilibrium maintaining structural constraints requisite for fitness balanced against the emergence of novel or improved biological functions. A protein multiple sequence alignment (MSA) organizes related homologues within a shared coordinate frame where conservation, covariation, and common domains can be conveniently modeled.^1–5^ As MSA methodology has advanced, faster homologue retrieval and embedding-based alignments provide sources of expanded information diversity for large-scale protein workflows.^6–9^

Protein generation and folding approaches generally utilize protein MSAs as fixed artifacts of homologue retrieval. Properties such as alignment depth or effective sequence count are referenced to denote the scale and diversity of the MSA. However, an MSA is of greater experimental value when evaluated holistically rather than categorized using singular descriptors. Alignments with identical depths might contain highly divergent evolutionary contexts; effective sequence count does not convey how information content is organized within sequences across an MSA. Perturbing MSA row composition could cause conservation, row similarity, and covariance to change together or to become entirely decoupled. Classical bioinformatic approaches such as direct-coupling and Potts-model approaches highlight that column-level covariation effectively encodes structural and functional constraints.^10–12^ Modern MSA-informed deep learning models utilize both row and column organization.^2,5^ These applications overlook an opportunity to tune an MSA’s row and column organization in a way that is independent from conservation and alignment depth.

Newer approaches augment or generate entirely synthetic MSAs. MSA-based protein language models can synthesize family-like sequences or even virtual MSAs to support cases where naturally occurring protein information is sparse or lacks sufficient heterogeneity.^6,13–15^ These results clearly demonstrate that evolutionary information can be manipulated to achieve a desired outcome. They do not, however, examine whether it is possible to steer protein model behavior based on controlling empirical MSA row composition.

In this study, evolutionary information is tuned through targeted row selection. Starting from single parental alignments across a diverse protein cohort, we use row selection to control lower-tail information content inside the projected coordinate frame of the reference protein. We also created a control matched-depth MSA with randomly selected rows for each information-tuned MSA. We then trained an MSA-informed masked diffusion language model (MDLM) that we used to generate de novo sequences for downstream analysis from each generated MSA.^16^ This experimental design isolates how evolutionary information shapes protein generation and structure prediction readouts while accounting for alignment depth.

A novel row-coherence metric empirically revealed a neighborhood at approximately 1 bit/site that separated low- and high-information regimes with distinct MSA topology and generated-sequence latent geometry properties. We show that MSA information content impacts MDLM decoder uncertainty, generated-sequence structural confidence and Chai-1 structure prediction accuracy in an independent JackHMMER folding benchmark.^1,17^ These results demonstrate that evolutionary information is not merely a passive result of homologue retrieval; it should be engineered and tuned.

## Results

Our experimental cohort consisted of 45 proteins representing diverse taxonomic backgrounds, functional classes, and sequence lengths from 27 species (Fig. 1a-c and Supplementary Data 1). We exploited this diversity to identify information-dependent patterns that were consistent across multiple protein families and organisms.

**Figure 1.**
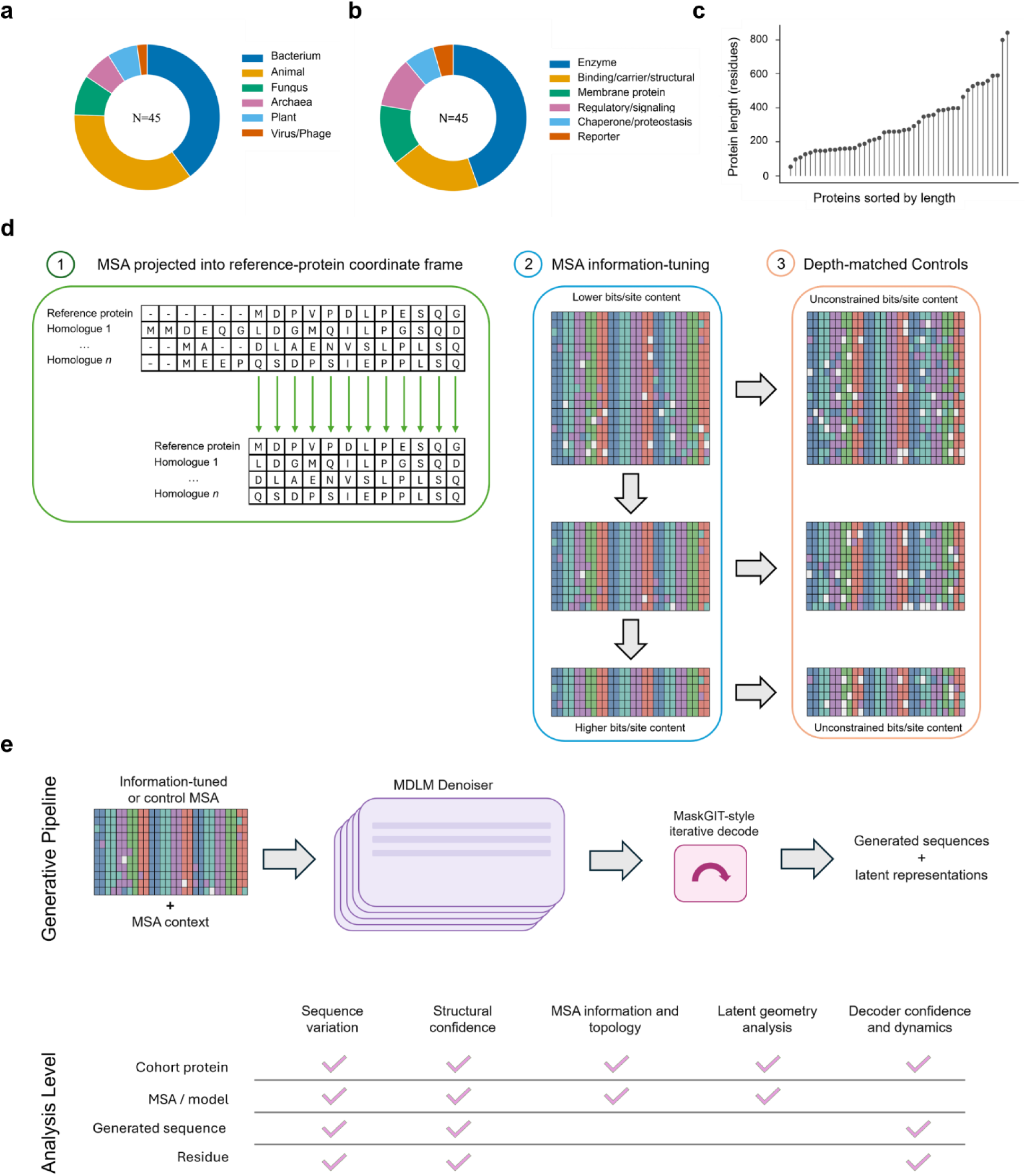
Cohort composition and MSA information-tuning workflow. **a**, Taxonomic composition of the 45-protein cohort. The cohort spans 27 species and includes bacterial (18/45), animal-derived (16/45), fungal, archaeal, plant, and virus/phage proteins. **b**, Functional composition of the cohort. Enzymes are the largest class (20/45), followed by binding, carrier, and structural proteins (9/45), membrane proteins (6/45), regulatory and signaling proteins (5/45), chaperone and proteostasis proteins (3/45), and reporters (2/45). **c**, Protein lengths across the study set, sorted by residue count. Lengths range from 54 to 843 residues, with a median of 262 residues. **d**, Workflow for tuning lower-tail multiple sequence alignment (MSA) information through row-selection. Parental MSAs are first projected through the reference-protein coordinate frame so that columns are fixed to the reference sequence. Row composition is adjusted toward predefined lower-tail bits/site targets using a row-coherence metric such that rows are dropped to greedily increase or decrease the metric. Finally, a depth-matched random-row control MSA is created for each information-targeted alignment. Information content is not targeted or constrained in the control MSAs. **e**, Generation and analyses workflow. Each information-targeted or matched-control MSA is used to train an MSA-informed masked diffusion language model (MDLM). Candidate de novo sequences are generated from the anchor sequence by masking non-gap positions and iteratively filling masked sites with a MaskGIT-style decoding schedule over twelve decode steps. Downstream categories of analysis performed in this paper are illustrated below the generative pipeline. The hierarchical levels at which analysis was performed in each category are noted by check marks. When appropriate, we report the results aggregated at the protein cohort level so that biological replicates are appropriately weighted.

MSA information was tuned by a row selection mechanism within a fixed alignment frame. We tuned MSAs to specific lower-tail information targets, such that an information-targeted MSA was created when the requested lower-tail target was achieved within tolerance. The lower-tail statistic was p10, defined as the 10th percentile of bits/site values across reference-projected columns with at least 50% occupancy. We tuned the MSAs across a grid of targets ranging from approximately 0.83 to 2.56 bits/site. For each generated information-targeted MSA, we also created a control MSA containing the same number of randomly selected rows. Comparing the targeted and control MSAs allowed us to isolate the impact of row coherence and information distribution while controlling for MSA depth (Fig. 1d).

Targeted and control MSAs always excluded insertions or deletions within the reference sequence’s match-state columns. Keeping only the columns that corresponded to residues in the reference sequence enforced a fixed coordinate frame such that information context was always comparable within the MSAs and de novo sequences generated for each cohort protein.

Each generated MSA was used to train an MSA-informed masked diffusion language model (MDLM) for de novo sequence generation (Fig. 1e). Because targeted and control MSAs were restricted to exactly the subset of columns containing reference protein residues, the de novo sequences produced by the MDLM models inherited this information-coordinate system. Deletions and insertions were not permitted in generated sequences. Therefore, sequence novelty is directly attributable to mutations rather than information-coordinate shifts.

### MSA-SNR identifies a row-coherence regime near 1 bit/site

We next asked whether information tuning via row selection would expose an information regime beyond the p10 statistic utilized for tuning.

Performing controlled row selection to realize lower-tail information targets impacted column-wise information content and altered the alignment composition of homologues and how homology was arranged in terms of pairwise-identity space. We measured this row structure using a novel metric, MSA-SNR, which we define as:

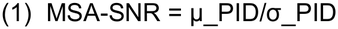

where μPID is the mean no-gap pairwise identity among retained rows and σPID is the corresponding standard deviation. We used MSA-SNR to select the rows that were removed from the parental MSA. Rows were removed by greedy-search to either increase or decrease MSA-SNR until a requested p10 value was achieved within tolerance. Downstream low- and high-information regimes were assigned independently using the MSA minimum bits/site value.

MSA-SNR did not change monotonically with p10 across the experimental cohort. Where the parental MSA’s pairwise identity was high, greedy-search row removal tended to produce positive correlations between MSA-SNR and p10 values. The opposite trend emerged where the parental MSA’s pairwise identity was low (Supplementary Fig. 1c). P07740 and P29972 provided illustrative examples (Supplementary Fig. 1a,b). Thus, the MSA-SNR metric behaves as a row-coherence metric rather than as a direct proxy for Shannon information.

We observed the minimum bits/site value where the highest MSA-SNR value was observed for each cohort protein. Across the experimental cohort, MSA-SNR peak locations clustered near 1 bit/site, with an unweighted mean of 0.79 ± 0.37 bits/site and a peak-Softmax-weighted mean of 0.96 ± 0.12 bits/site (Fig. 2e,f). We would note that because row selection realized lower-tail p10, this boundary was not directly imposed by the tuning process. Across cohort MSAs, the p10 and minimum bits/site did not always move together as p10 effects can appear compressed at the alignment minimum (Supplementary Fig. 1d). Additionally, we consider the 1-bit/site value an empirical reference point for this cohort rather than a universal threshold. When we assessed the stability of these comparisons at minimum-information thresholds of 0.8, 0.9, 0.972, 1.0, 1.1, and 1.2 bits/site, the direction of 15 of the 17 features shown in Figs. 3 and 4 remained consistent across all thresholds (Supplementary Data 6). For this reason, comparisons above and below the operational threshold should be read as contrasts across a local information neighborhood where differences exist as a continuum rather than a sharp boundary.

**Figure 2.**
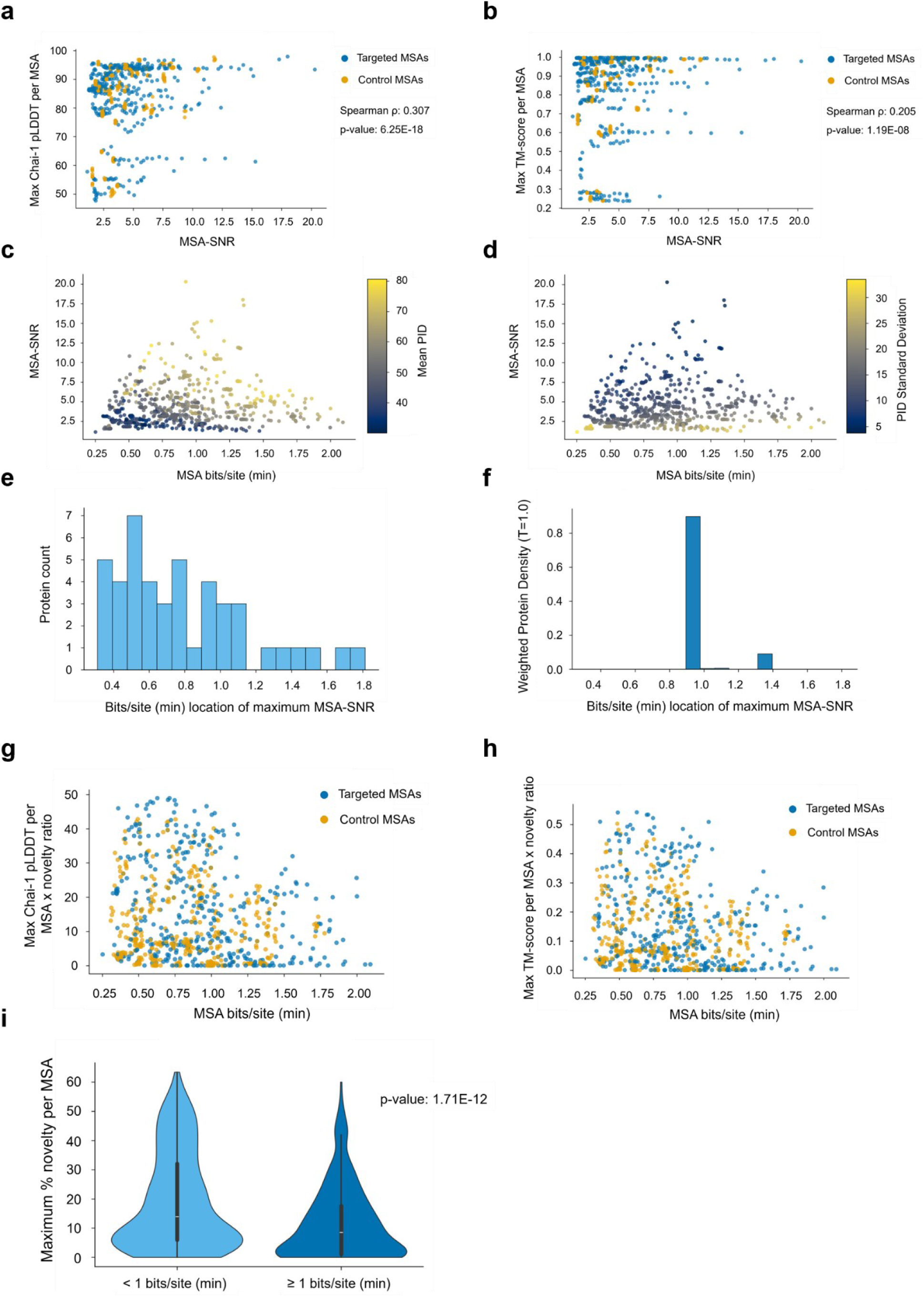
MSA-SNR identifies a row-coherence regime near 1 bit/site. **a**, Maximum mean Chai-1 pLDDT across generated sequences from each MSA, plotted against MSA-SNR (Spearman ρ = 0.307, p = 6.25 × 10^-18, n = 757). **b**, Maximum mean TM-score to the Chai-1-predicted anchor structure across generated sequences from each MSA, plotted against MSA-SNR (Spearman ρ = 0.205, p = 1.19 × 10^-8, n = 757). **c**, MSA-SNR versus minimum bits/site per MSA, colored by mean no-gap pairwise identity. **d**, MSA-SNR versus minimum bits/site per MSA, colored by no-gap pairwise-identity dispersion. **e**, Distribution of the minimum bits/site values at which each cohort protein reaches its maximum observed MSA-SNR. The unweighted mean peak location is 0.79 ± 0.37 bits/site, and 73.3% of proteins peak below the operational threshold of 0.972 bits/site. **f**, Softmax-weighted distribution of per-protein MSA-SNR peak locations using each protein’s maximum observed MSA-SNR value as the weight source, with τ = 1.0. The weighted mean peak location is 0.96 ± 0.12 bits/site. **g**, Maximum pLDDT-based novelty-frontier score plotted against minimum bits/site per MSA. For each de novo sequence, we compute mean pLDDT × novelty expressed as a ratio. The maximum value from each MSA is displayed. **h**, Maximum TM-score-based novelty-frontier score plotted against minimum bits/site per MSA. For each de novo sequence, we compute TM-score × novelty expressed as a ratio. The maximum value from each MSA is displayed. **i**, Maximum generated-sequence novelty per MSA below and at or above the operational threshold of 0.972 bits/site. Novelty is defined as 100 minus maximum percentage sequence identity to the BLAST-derived sequences used for MSA construction and is higher below the operational threshold (19.33% ± 16.67%, n = 482, versus 11.75% ± 12.55%, n = 292; Welch two-sided p = 1.71 × 10^-12).

**Figure 3.**
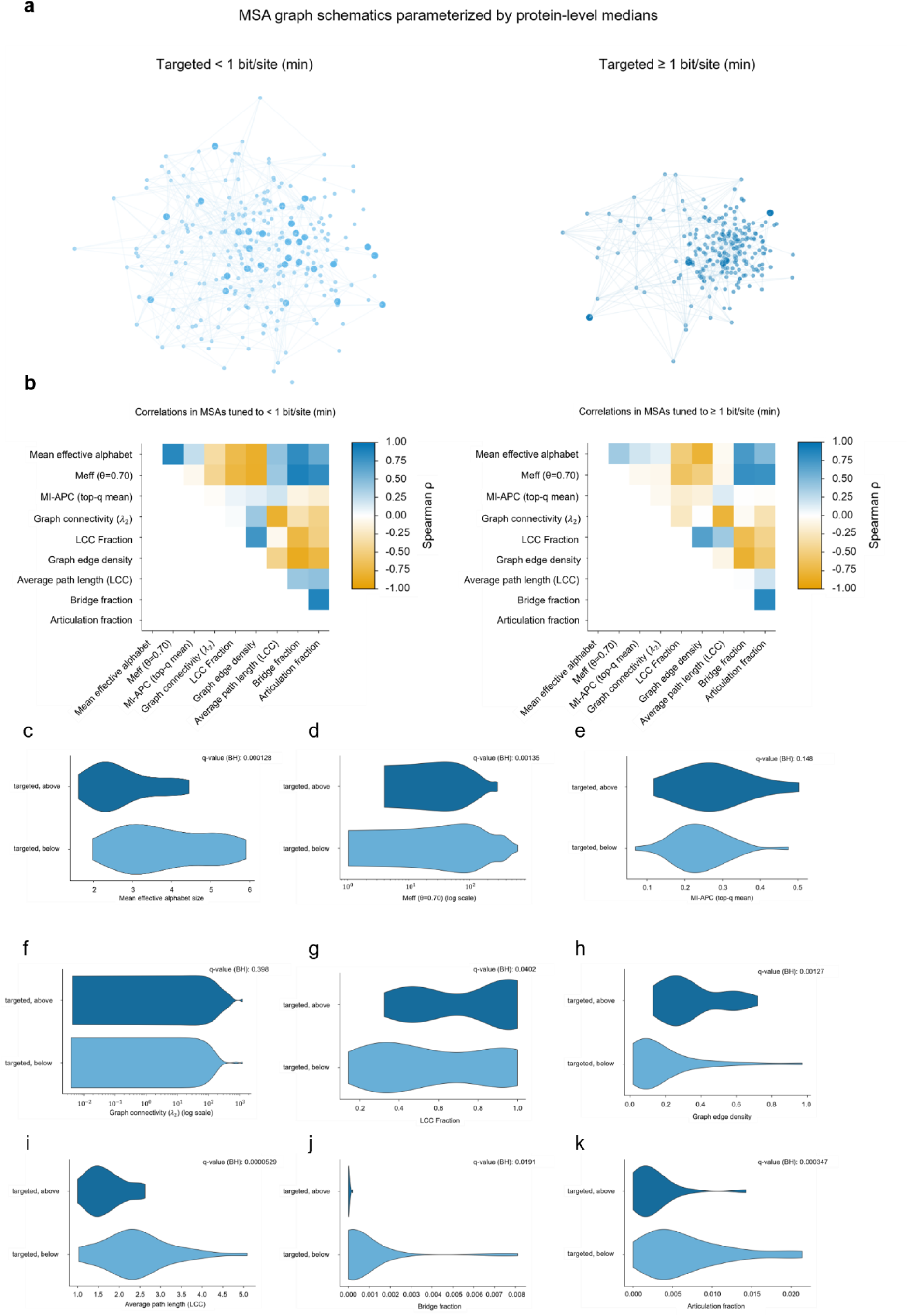
MSA row-similarity topology distinguishes the MSA-SNR-centered regime. **a**, Parameterized MSA graph schematics generated from protein-level median graph summaries. **b**, Upper-triangle Spearman correlation heatmaps for protein-level mean MSA features. Mean absolute correlation is 0.483 below the reference point and 0.351 at or above it. Median absolute correlation is 0.439 and 0.265. **c**, Mean effective alphabet size is greater below the operational threshold than above it (3.821 ± 1.151 versus 2.763 ± 0.840; Welch two-sided p = 6.39 × 10^-5, BH q = 1.28 × 10^-4). **d**, Meff at θ = 0.70 is significantly higher below the reference point (202.06 ± 168.96 versus 87.08 ± 85.88; Welch two-sided p = 6.77 × 10^-4, q = 0.00135). **e**, Top-quantile APC-corrected mutual information trends higher above the reference point, but does not significantly differ after correction (0.241 ± 0.0722 versus 0.281 ± 0.100; Welch two-sided p = 0.0828, q = 0.148). **f**, MSA graph connectivity λ₂ trends higher above the reference point, but does not significantly differ after correction (68.244 ± 236.132 versus 152.376 ± 270.713; p = 0.199, q = 0.398). **g**, LCC fraction is significantly lower below the reference point (0.573 ± 0.301 versus 0.739 ± 0.255; p= 0.0201, q = 0.0402). **h**, Edge density is significantly lower below the reference point (0.175 ± 0.203 versus 0.342 ± 0.169; p= 6.33 × 10^-4, q = 0.00127). **i**, Average path length within the LCC is significantly higher below the reference point (2.415 ± 0.843 versus 1.669 ± 0.470; p = 2.65 × 10^-5, q = 5.29 × 10^-5). **j**, Bridge fraction is significantly higher below the reference point (7.48 × 10^-4 ± 0.00171 versus 3.36× 10^-5 ± 4.55 × 10^-5; p = 0.0140, q = 0.0191). **k**, Articulation fraction is significantly higher below the reference point (0.00669 ± 0.00544 versus 0.00250 ± 0.00291; p = 1.74 × 10^-4, q = 3.47 × 10^-4).

**Figure 4.**
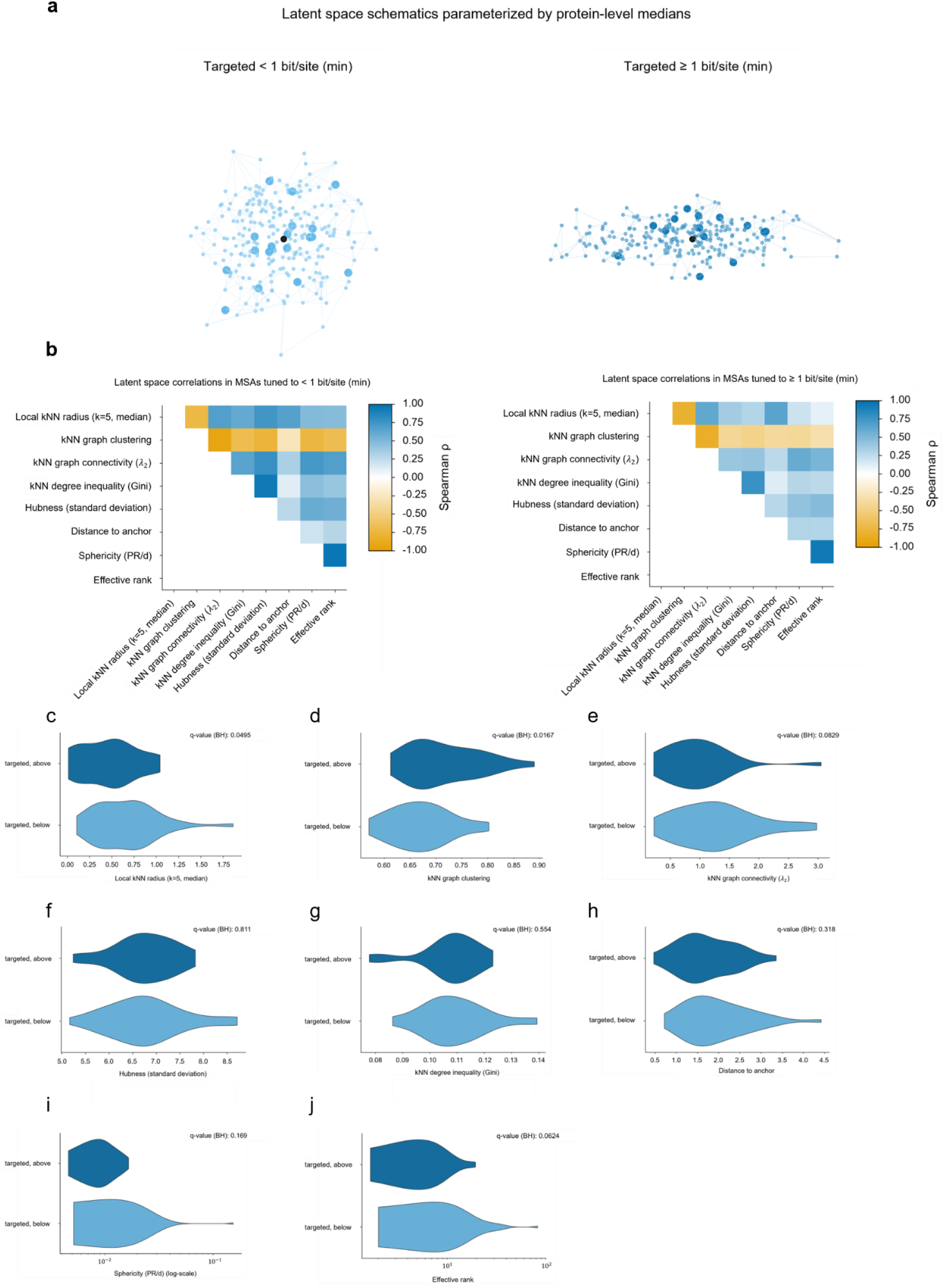
Lower-information MSAs broaden generated-sequence latent neighborhoods. **a**, Parameterized latent-space schematics generated from protein-level median latent-geometry summaries. The anchor/reference latent is displayed as a black dot. **b**, Upper-triangle Spearman correlation heatmaps for protein-level mean latent-geometry features. Mean absolute correlation is 0.582 below the reference point and 0.435 at or above it. Median absolute correlation is 0.653 and 0.400. **c**, Local kNN median radius (k = 5) is significantly higher below the reference point (0.656 ± 0.347 versus 0.465 ± 0.316; Welch two-sided p = 0.0248, Benjamini-Hochberg q = 0.0495). **d**, Latent kNN graph clustering is significantly lower below the reference point (0.671 ± 0.0578 versus 0.715 ± 0.0687; p = 0.00837, q = 0.0167). **e**, Latent kNN graph connectivity λ₂ trends higher below the reference point but does not differ after correction (1.284 ± 0.662 versus 0.966 ± 0.565; p = 0.0415, q = 0.0829). **f**, Latent hubness, measured as the standard deviation of directed kNN neighbor-occurrence counts, is similar across regimes (6.788 ± 0.809 versus 6.745 ± 0.642; p = 0.811, q = 0.811). **g**, kNN degree inequality changes little between regimes (0.109 ± 0.0119 versus 0.107 ± 0.0110; p = 0.503, q = 0.554). **h**, Distance to the anchor/reference latent in the original latent coordinate space trends higher below the reference point, but is not significant (1.953 ± 0.763 versus 1.713 ± 0.668; p = 0.183, q = 0.318). **i**, Sphericity, defined as participation ratio divided by latent dimensionality, trends higher below the reference point, but is not significant after correction (0.0165 ± 0.0238 versus 0.00955 ± 0.00332; p = 0.0843, q = 0.169). **j**, Effective rank trends higher below the reference point but does not differ after correction (12.398 ± 14.364 versus 6.888 ± 4.409; p = 0.0312, q = 0.0624).

We used Chai-1 for structural analysis of the generated de novo sequences, as the Chai-1 model allowed us to generate structural predictions in an MSA-free manner. The row-coherence metric, MSA-SNR, was positively correlated with the maximum mean pLDDT of the sequences generated from each MSA as well as the maximum TM-score of generated sequences to the Chai-1 predicted reference structure (Fig. 2a,b). The random-row depth-matched controls showed the same positive correlations even though their rows were not selected using MSA-SNR or even with any information content objective. The same correlations were weaker between Chai-1 structural readouts and Meff or alignment depth (Supplementary Fig. 2a-d).

Thus, the MSA-SNR metric connected row coherence to structural confidence and sequence exploration. MSAs where the minimum bits/site value fell below the operational threshold tended to generate de novo sequences with higher novelty, while MSAs at or above this threshold produced sequences with lower diversity. The confidence-novelty tradeoff is centered near the per-protein MSA-SNR peak at approximately 1 bit/site (Fig. 2g-i). When we evaluated MSA-SNR against the MSA-level structural readouts, we note that the random-row control MSAs often exceeded the correlations shown in the information-targeted MSAs. These relationships also persisted after adjusting for the effects of Meff or alignment depth, although Meff did retain weak residual associations within a partial correlation analysis. Additionally, protein-block subsampling confirmed that these relationships were not driven by the presence of a small subset of the protein cohort (Supplementary Data 6).

Further component analysis of the MSA-SNR metric illustrated that pairwise dispersion is the main driving component behind the MSA-SNR peak. When we examined the numerator and denominator of the MSA-SNR metric separately, we observed that near the empirical reference point alignments with high MSA-SNR maintained higher mean PID with lower PID dispersion (Fig. 2c,d). When we performed sensitivity analysis to test whether this result relied on a specific component of the MSA-SNR ratio, effective sequence count, or alignment depth, we found that negative pairwise-identity dispersion most closely reproduced the primary MSA-SNR peak location pattern (Supplementary Data 6). This indicated that reduced dispersion across retained rows, not mean pairwise identity, was the primary factor. The fact that these relationships persisted after adjustment for Meff and retained depth (Supplementary Data 6) signaled that how information is organized across rows in the neighborhood around the 1-bit/site minimum-information reference point was the most salient factor, not how many sequences are present or how diverse they are.

These analyses identified a reference point near 1 bit/site that characterizes an intermediate MSA-information neighborhood. Within this neighborhood, retained rows remain coherent enough to maintain key structural constraints yet downstream generative models still have room to explore.

### MSA row-similarity topology distinguishes the MSA-SNR-centered regime

We next tested whether the MSA-SNR-established reference point reflected a significant change in how rows were organized within MSAs. We represented each MSA as a row-similarity graph such that rows were connected when no-gap pairwise identity was ≥ 0.70. All graph metrics were computed as protein-level averages, which were used for comparing between the below-threshold and above-threshold information-tuned MSAs.

The comparisons shown in Fig. 3 use unpaired Welch tests as not every protein was represented on both sides of the reference point. Metric definitions, regime-test summaries and feature-correlation summaries for the MSA-topology and latent-geometry analyses are provided in Supplementary Data 3. Fig. 3a is a parameterized schematic illustrating protein-level median graph values for targeted MSAs below and above the 1-bit/site minimum reference point.

Lower-information MSAs reflected greater sequence diversity as effective alphabet size and Meff were both higher when the MSA bits/site minimum was below the 1-bit/site reference point (Fig. 3c,d). However, top-quantile APC-corrected mutual information was not significantly different (Fig. 3e). We also observed that protein-level MSA metrics were more strongly coupled below the threshold than above it (Fig. 3b).

Analysis of graph topology features showed the strongest separation between below- and above-threshold MSAs. Largest connected components (LCC) fractions were significantly smaller in the low-information regime. Edge densities were significantly lower and LCC path lengths were significantly longer in low-information MSAs. These features describe MSA row-similarity graphs that tended to be more fragmented and less compact below the 1-bit/site minimum-information threshold (Fig. 3g-i).

The graph fragility metrics, bridge and articulation fractions, were also higher in lower-information MSAs, indicating that the nodes in these graphs were more susceptible to disconnection due to greater reliance on narrow connections between neighborhoods (Fig. 3j,k). Graph connectivity was increased in above-threshold MSAs but this difference was not significant (Fig. 3f).

Below the empirical 1-bit/site reference point, information-tuned MSAs demonstrated higher sequence diversity, but this diversity was represented through more fragmented and bottlenecked MSA similarity graphs.

### Lower-information MSAs broaden generated-sequence latent neighborhoods

We next asked whether we could observe similar effects in the latent representations of MDLM-generated sequences. Fig. 4a displays a parameterized schematic illustrating protein-level latent geometry median values, while Fig. 4c-j quantifies latent geometry metrics for sequences generated from information-targeted MSAs. As in Fig. 3, we observed stronger correlation between latent features from information-targeted MSAs below the 1-bit/site threshold compared to those above the threshold (Fig. 4b).

We observed that the strongest contrast between below- and above-threshold MSAs when measuring the local kNN radius and graph clustering of generated-sequence latent representations (Fig. 4c,d). The latent representations downstream of below-threshold MSAs displayed significantly larger local kNN radii and significantly lower latent graph clustering compared to latent representations from above-threshold MSAs. These observations describe latent representations generated from below-threshold MSAs as being composed of broader, less organized local neighborhoods. This finding mirrored the less compact nature of the below-threshold MSA similarity graphs.

Differences between the other latent geometry descriptors shown in Fig. 4e-j were not significant within our experimental cohort. Below-threshold MSAs all had higher average connectivity, distance to anchor (where anchor is the reference protein latent representation), sphericity, and effective rank values. Hubness and degree inequality were nearly identical across all information-targeted MSAs.

Because not every parental MSA in the experimental cohort could be tuned to both sides of the 1-bit/site threshold, we performed a paired sensitivity analysis using only the proteins that could produce information-targeted MSAs in both regimes. Across these 20 proteins, Fig. 3 metrics preserved their direction. Meff, edge density, average path length, and articulation fraction all remained significant across the paired comparison. The latent geometry metrics described in Fig. 4 all maintained the same trends and direction of effects, but no individual latent metric was significant in the smaller paired subset (Supplementary Data 6).

These results demonstrate that effects observed in the lower-information regime extend beyond the alignments into the latent representations of generated sequences. The combined results suggest that the downstream impacts of tuning an MSA to below the 1-bit/site minimum-information threshold tend to be localized rather than global and these changes result in broader neighborhood radii with less clustering.

### Targeted low-information MSAs concentrate mutations and decoder uncertainty

We next asked how protein-level MSA analysis would relate to properties of generated de novo sequences such as mutation rates, MDLM decoder entropy, and downstream Chai-1 structural readouts. In these analyses, we considered a residue to be mutated if the residue was edited relative to the corresponding residue in the reference sequence since MDLM models were conditioned with a corrupted reference sequence to generate each de novo sequence. The profiles in Fig. 5 use protein-normalized summaries so that residue-level analyses provided equal weight to each cohort protein regardless of sequence length or the number of de novo sequences generated.

**Figure 5.**
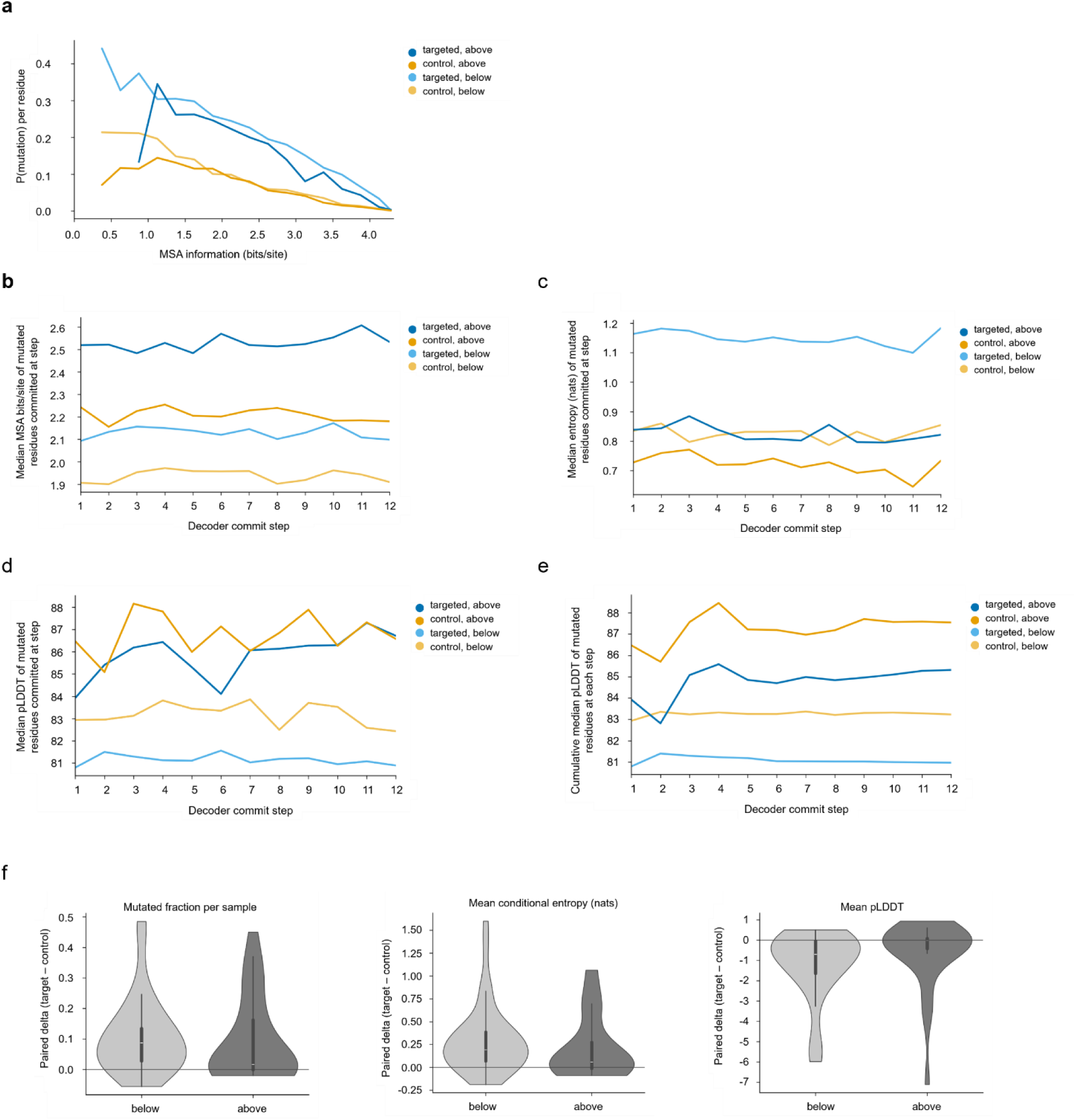
Targeted low-information MSAs concentrate mutations and decoder uncertainty. **a**, Mutation probability as a function of local MSA bits/site. Residues were binned by local bits/site within each protein and condition, and plotted profiles show medians across contributing proteins. **b**, Protein-median MSA bits/site of mutated residues committed at each decode step. **c**, Protein-median conditioned decoder entropy, in nats, of mutated residues committed at each decode step. **d**, Protein-median Chai-1 pLDDT of mutated residues committed at each decode step. **e**, Cumulative protein-median Chai-1 pLDDT of mutated residues committed at or before each decode step. **f**, Paired information-targeted minus matched-control deltas for mutated fraction, mean conditioned decoder entropy, and mean Chai-1 pLDDT, stratified by below- and above-threshold targeted MSAs. Positive values indicate higher values in the information-targeted condition; n = 38 lower-information and n = 27 higher-information protein-level pairs.

The information content at column positions demonstrated a clear relationship between mutational probabilities and the information content of corresponding MSA positions (Fig. 5a). Residues predicted from lower-information columns were more likely to be mutated. This probability decreased steadily as the bits/site content of columns increased. Information-targeted MSAs showed higher mutation probabilities compared to matched-depth control MSAs at positions containing comparable quantities of information with the greatest separation in the lower-information regime. This result demonstrates that sequence novelty generally arises from low-information positions in MSAs rather than as an artifact of MDLM models, which illustrates the importance of information organization within MSAs.

We observed a correlative trend across decoding steps. Mutations from models trained with targeted below-threshold MSAs occurred in lower-information positions, and MDLM models trained with targeted low-information MSAs demonstrated the highest entropy when generating these mutations (Fig. 5b,c). Low-information MSAs tended to have increased model uncertainty and sequence exploration with the largest shift occurring in the targeted below-threshold regime. Additionally, when corrupted sequence states were evaluated with and without an MSA context array, we observed that MSA context impacted MDLM confidence most strongly at low-information positions (Supplementary Fig. 3a,b).

Sequences generated from below-threshold information-targeted MSAs showed lower Chai-1 pLDDT at mutated residues compared to matched controls. This confidence decrease was consistent across individual decoding steps and cumulatively across successive steps (Fig. 5d,e). Additionally, model uncertainty varied with Chai-1 pLDDT more closely in the information-targeted MSAs than in the matched controls (Supplementary Fig. 3e). Analysis of model uncertainty-confidence agreement suggested that these differences were not simply due to significant systematic miscalibrations across the four conditions (Supplementary Fig. 3c,d).

To assess whether these trends were consistent at the protein level, we compared information-targeted MSAs with their paired matched-depth control MSAs. We first computed cohort-protein averages and then evaluated differences between targeted-minus-control averages below and above the 1-bit/site minimum-information threshold (Fig. 5f and Supplementary Data 4). Information-targeted conditions showed higher rates of mutation and decoder uncertainty with lower Chai-1 pLDDT compared to their matched controls. These differences were larger in the below-threshold information-targeted MSAs than in their above-threshold counterparts. Collectively, these results show that targeting MSA information shifts mutation rates, model confidence, and downstream structural confidence, and that these effects are organized through MSA information content rather than driven by alignment depth.

### Local MSA information tracks residue-level structural confidence

We next asked how local MSA information co-varied with residue-level MDLM uncertainty and Chai-1 structural confidence readouts. For each sequence, we compared local MSA bits/site with MSA-conditioned decoder entropy, per-residue Chai-1 pLDDT and mean per-residue Chai-1 PAE. Fig. 6a highlights two example sequences generated from P07740. The plots generally show that local MSA information tends to positively correlate with Chai-1 pLDDT and negatively correlate with MDLM entropy and Chai-1 PAE.

**Figure 6.**
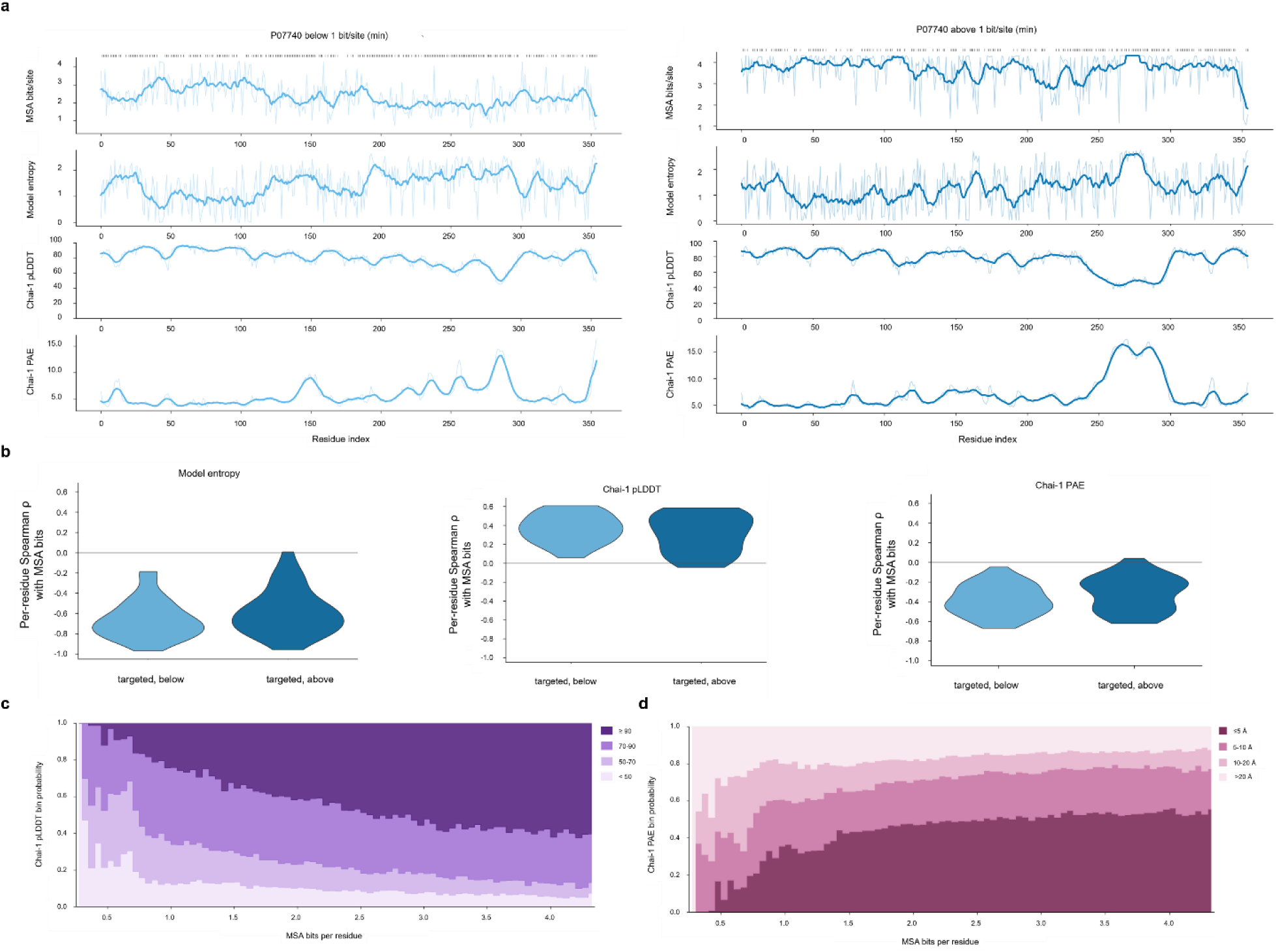
Local MSA information tracks residue-level structural confidence. **a**, Per-residue traces for P07740 from information-targeted MSAs below and above the 1-bit/site threshold. Rug marks indicate positions where a mutation was made relative to the reference sequence and the trend lines illustrate rolling nine-residue window averages. In the below-threshold example, local MSA information shows stronger residue-level associations with decoder entropy (ρ = - 0.729), pLDDT (ρ = 0.387), and mean PAE (ρ = -0.421) than in the above-threshold example. The corresponding higher-information correlations are entropy ρ = -0.470, pLDDT ρ = -0.116, and mean PAE ρ = -0.024. **b**, Protein-level local information flow, summarized as residue-level Spearman correlations between local MSA bits/site and each downstream per-residue metric after aggregating sample-level correlations to protein-level condition summaries. For information-targeted MSAs, mean ± s.d. Spearman ρ values are decoder entropy, -0.657 ± 0.190 for samples below the 1-bit/site threshold and -0.584 ± 0.226 in above-threshold samples; Chai-1 pLDDT, 0.365 ± 0.153 and 0.308 ± 0.200; and mean PAE, -0.385 ± 0.155 and -0.319 ± 0.178. **c**, Protein-balanced probabilities of Chai-1 pLDDT confidence bands across local MSA bits/site bins across generated samples. Residues are assigned to fixed-width local bits/site bins and summarized over pLDDT bands <50, 50-70, 70-90, and ≥90. **d**, Protein-balanced probabilities of per-residue mean PAE bands across the same local MSA bits/site bins across generated samples. Residues are summarized over PAE bands ≤5, 5-10, 10-20, and >20.

Analysis across the protein cohort showed that the same residue-level coupling was present across the entire dataset. We initially computed these correlations at the residue level for each generated protein. The results were then balanced at the protein level to equally weight cohort proteins regardless of their length or number of sequences generated. As in the P07740 example, local MSA information negatively correlated with MDLM entropy and mean per-residue Chai-1 PAE but positively correlated with per-residue Chai-1 pLDDT. These correlations were strongest in the below-threshold information-targeted MSAs, mirroring the earlier MSA topology and latent geometry metric results (Fig. 3b, Fig. 4b).

Confidence-band plots of Chai-1 structural confidence probabilities across local bits/site bins showed the same shift at the local residue level. Bins representing protein-balanced local information illustrate that Chai-1 pLDDT is lower and mean per-residue PAE is higher for lower-information positions (Fig. 6c,d). Bins with higher bits/site content displayed higher pLDDT and lower mean per-residue PAE. We also note that pLDDT and PAE both show large shifts in structural confidence probabilities as information content decreases below the 1-bit/site reference threshold. The decline in structural confidence below 1 bit/site was even more noticeable when we considered only mutated residues (Supplementary Fig. 4b,c). For the mutated residue plots in Supplementary Fig. 4b,c, we provided separate plots of the below-threshold and above-threshold targeted conditions to illustrate that these lower confidence bands simply do not occur in the above-threshold condition when there are no positions with less than 1 bit/site in the MSAs.

To qualitatively test information flow potential from MSA to generated sequence structure, we performed mediation-style screening to identify putative covariance chains (Supplementary Fig. 4a). Our screen assessed serial chains of covariance from MSA-level information content to MSA and latent features we analyzed in Figs. 3 and 4 to Chai-1 structural readouts (per-residue PAE and pLDDT). This screen was not meant to prove causal mediation for specific chains. We use it here to demonstrate that there were many more putative, reproducible MSA-to-latent-to-Chai-1 PAE covariance chains in targeted below-threshold MSAs than in targeted above-threshold MSAs.

Our analyses demonstrate that, at residue scale, there is a model confidence drop at low-information positions. These low-information positions demonstrate higher MDLM decoder uncertainty and lower Chai-1 structural confidence, with mutated residues reflecting this trend most strongly.

### MSA tuning around the 1-bit/site reference point identifies confidence-gated structure-prediction gains

In the preceding analyses, our approach was to tune the lower-tail information content of MSAs and then to use these MSAs to train and condition protein generative models. Chai-1 structure prediction was performed without MSAs to avoid introducing a confounding variable. We next asked whether this approach could be practically engineered for MSA-informed Chai-1 structure prediction.

In MSA, latent, and local information flow analyses, we observed that MSAs with positions below the operational threshold exhibited stronger information coupling. For this reason, we anticipated that low-information positions might contain covariance patterns that are useful for identifying structural constraints. To test this hypothesis, we generated JackHMMER alignments for a benchmark dataset of diverse, single polymer chain structures deposited after 2022. We tuned the row composition of these JackHMMER alignments to control the fraction of reference protein-projected columns where information fell below the approximately 1-bit/site reference point. Chai-1 predictions were performed using these tuned MSAs as input, and we compared structural accuracy relative to inference using parental JackHMMER MSAs (Fig. 7a,b and Supplementary Data 5).

**Figure 7.**
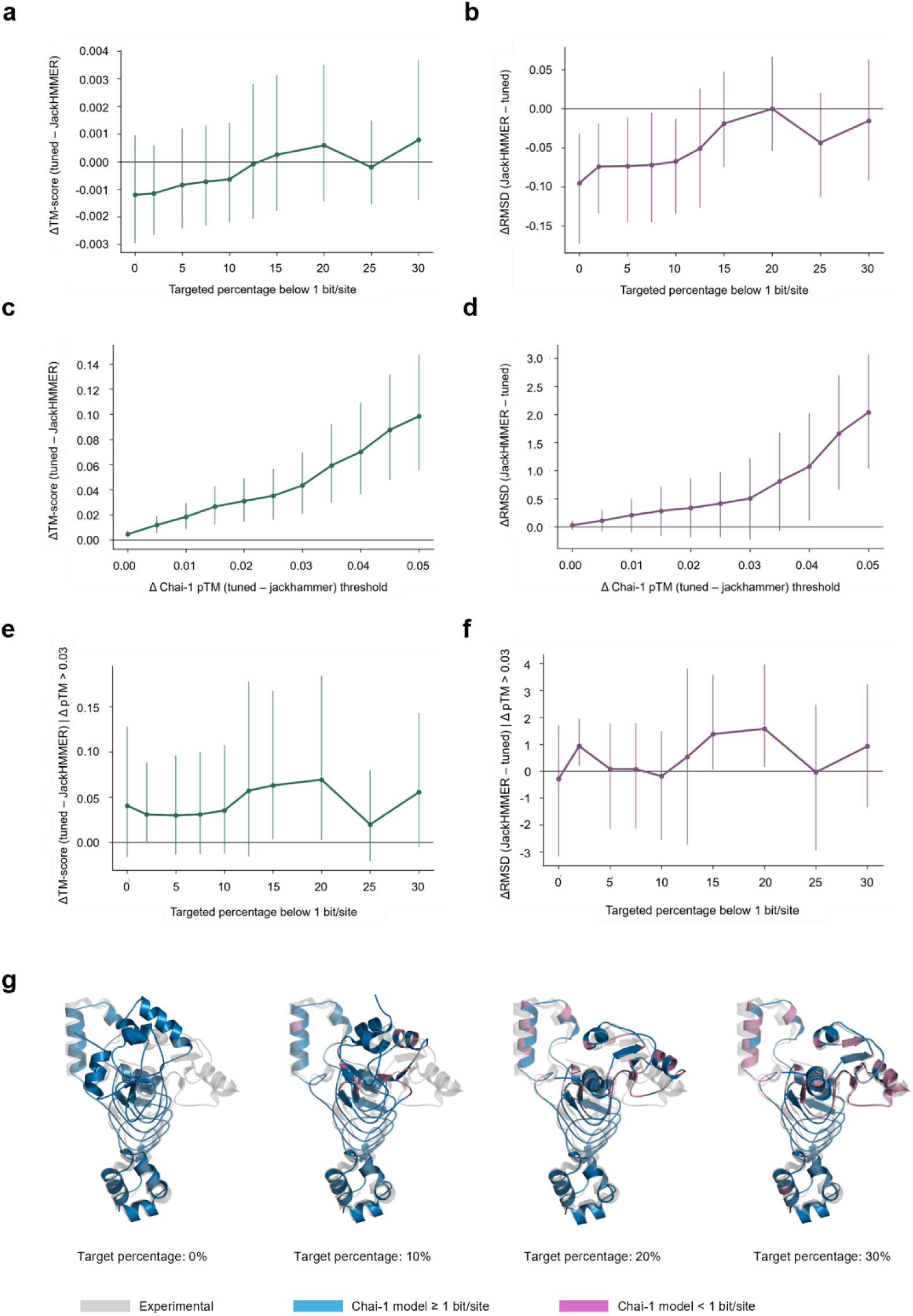
MSA tuning around the 1-bit/site reference point identifies confidence-gated structure-prediction gains. **a**, Mean ΔTM-score as a function of the fraction of columns below the 1-bit/site reference point. Mean effects across the full matched set remain close to zero. **b**, Mean ΔRMSD across the same target range. Global RMSD penalties at the lowest fractions weaken as the below-reference fraction increases. **c**, Mean ΔTM-score as a function of the ΔpTM gate threshold after pooling matched pairs across target percentages. **d**, Mean ΔRMSD as a function of the same ΔpTM gate threshold for pooled comparisons. **e**, Mean ΔTM-score as a function of target fraction after applying the primary ΔpTM > 0.03 confidence gate. Gains are enriched at intermediate fractions below the reference point. **f**, Mean ΔRMSD under the same ΔpTM > 0.03 gating criterion. Improvement bands are most apparent at intermediate target fractions. **g**, Structural overlay for PDB 7UUJ comparing tuned-MSA predictions against the experimental structure. Residues in the tuned prediction are colored by whether the corresponding MSA column falls below the 1-bit/site reference point. Protein models generated from below-threshold fractions 0, 10, 20, and 30% are displayed as a progression.

Many of our parental MSAs were unable to produce tuned MSAs for each requested below-threshold fraction target. To avoid survivorship biases, we only considered the 441 structures where we had matched tuned MSAs and structural analysis for every requested below-threshold fraction target.

Across the matched benchmark structures, the average effects of controlling the fraction of columns below the information threshold were small. The mean ΔTM-score remained near zero across the requested fractions, although ΔRMSD showed that tuned MSAs with low fractions of columns below the information threshold performed significantly worse than the matched JackHMMER parental MSAs.

In cases where the Chai-1 pTM confidence was higher when making predictions with a tuned MSA compared to the JackHMMER MSA, we saw that this enriched for predicted structures displaying larger gains in TM-score and RMSD (Fig. 7c,d). We observed that the strongest improvements in structure prediction accuracy occurred at intermediate target fractions where 15-20% of columns were below the empirical information threshold when we used a primary gating strategy of isolating cases where tuned MSAs increased the Chai-1 pTM by more than 0.03 relative to the parental JackHMMER MSA (Fig. 7e,f).

Analysis of MSA alignment depth and Meff (θ = 0.70) in the tuned MSAs showed that both metrics increased monotonically as the fraction of columns below the approximately 1-bit/site threshold increased (Supplementary Fig. 5a,b). Both depth and Meff continued to rise beyond the 15-20% range while the confidence-gated structural accuracy decreased at higher fractions. This discrepancy does not mean that alignment depth and effective sequence count were not contributing factors, but it militates in support of the conclusion that the observed improvement in folding accuracy is driven by more than these two metrics alone.

The 7UUJ overlay in Fig. 7g highlights an example where the placement of low-information columns correlated with improvements in structure accuracy. The residues informed by MSA positions above the threshold and below the threshold are colored separately and overlaid on the experimental structure. As the fraction of below-threshold MSA columns increases, low-information positions concentrate where the structural accuracy is most improved. This example illustrates that the placement of the below-threshold columns can be just as important as their quantity.

We performed local Cα-error analyses within 11-residue windows across the benchmark dataset. We classified each 11-residue window by whether its mean bits/site lay above or below the 0.972-bit/site threshold in the parental JackHMMER MSA and tuned MSAs. We assessed the impact of the four possible local transitions, with respect to the information reference point, between each JackHMMER parental window and the corresponding tuned MSA window (above→above, above→below, below→below, below→above). The observed trend is that the largest structural accuracy improvements are concentrated in the above→below transition class (Supplementary Fig. 5c). Gating on examples where tuning increased Chai-1 pTM by more than 0.03 makes the trend even more apparent (Supplementary Fig. 5d). Within the benchmark dataset, structural accuracy outcomes depended not just on the below-threshold column budget, but on where in the MSA that budget was spent relative to the parental alignment.

These analyses confirm that the empirical information reference point we identified using de novo sequence generation can also organize outcomes in structural prediction tasks. We hypothesize that we saw modest accuracy changes across the benchmark dataset because the Chai-1 structure prediction method is not reliant on an MSA input for accurate inference. However, even in this context, we observed significant confidence-gated improvements at intermediate fractions of below-threshold positions, and the structural accuracy gains were predominantly localized within a specific information-state transition.

## Discussion

Generative and protein structure-prediction models frequently use MSA inputs as the fixed endpoint of a homologue retrieval task. In this paper, we show that the lower-tail information content of an MSA can be tuned through row selection in the context of a fixed information coordinate frame. This process revealed an intermediate information point that organized de novo generated sequence novelty and structural confidence readouts.

Within our generative experiments, the raw or effective number of sequences an MSA contained did not fully explain downstream outcomes. MSA depth alone was not predictive of structural outcomes, while Meff displayed only a partial effect. MSA-SNR, a novel row-coherence metric, was much more predictive of Chai-1 pLDDT and the TM-scores between de novo and reference proteins. The protein-level MSA-SNR peak defined the location of a structural confidence-novelty frontier. Below this point, generated sequence novelty becomes higher, but this is at the expense of decoder uncertainty and downstream structural confidence readouts. These effects were observed at both the MSA-level and the residue-level and are explained by protein-level analyses of MSA topology, generated-sequence latent properties, and decoder entropy.

These results were not explained by the fact that MSA-SNR was used for row-selection in the information-targeted MSAs. The relationship between MSA-SNR and novelty and structure confidence results persisted even when MSA rows were selected randomly. In some comparisons, correlations were even stronger within the random-row control MSAs than with information-targeted MSAs. Additionally, partial correlation analysis showed that the MSA-SNR metric was still associated with downstream readouts even after accounting for the contributions of the Meff and alignment depth metrics.

Across the 45-protein cohort, protein MSAs on either side of the 1-bit/site threshold showed significant differences in terms of row-topology, generated-sequence latent properties, mutation probabilities, and decoder uncertainty. When we explored other proximate thresholds, the results were largely directionally and statistically consistent. Thus, we describe the 1-bit/site reference point as a local information neighborhood where evolutionary constraints are relaxed enough to support novelty yet coupled enough to maintain fold-relevant signal.

These outcomes are consistent with prior results showing that naturally occurring protein folds are specified by well-defined minimum information constraints.^18^ By treating MSA information as a tunable parameter, we show that this principle can be extended to MSA-informed model behavior. Outcomes across both generative and structural prediction experiments depended on more than the amount of information contained in an MSA; outcomes depended on how that information is organized.

The 1-bit/site threshold could also be used to steer Chai-1 confidence-gated accuracy across an independent JackHMMER benchmark dataset. These results contextualized a key implication from the generative experiments. While MSAs with 15 and 20% fractions of columns below the threshold demonstrated improved accuracy after confidence-gating, local transition analysis clarified that these improvements were linked to the context in which the lower-information columns were placed. For this reason, we expect that MSA-tuning strategies tend to work best when they use protein structure to guide the precise placement of MSA information.

A similar principle is likely to be useful for generative design tasks. From our residue-level analyses, we infer that an optimal MSA information-tuning strategy must balance task-oriented information placement against the tradeoff between decoder uncertainty and structural confidence. Motifs that must be protected or maintained should utilize higher information content at those positions, while targeted exploration can be encouraged by relaxing information at locations where novelty is useful.

Our analysis also suggests a new class of protein model where an MSA is not treated as a singular input, but rather as the precursor to a panel of information-controlled MSAs. This class of protein model would not passively utilize a fixed MSA but rather create or select an optimal MSA that is best suited for the design or prediction task at hand. In this approach, evolutionary information is more than a mere input; it becomes an integral part of the inference process.

We acknowledge that these results are specific to the datasets, alignment methodology, and model stack used in this paper. The tuning process described herein is limited by the fact that it controls information indirectly through row selection, rather than by specifically targeting where information is placed in the MSA. The Fig. 7 structure prediction improvements were only significant after confidence gating, and the structural readouts in this paper were derived solely from Chai-1 inference rather than from experimental methods.

These results elucidate several clear relationships between the information content of MSAs and the downstream behavior of models that use them. They also bridge a conceptual gap between classical direct-coupling methods and how MSA information is utilized in modern protein deep learning models. Our results clearly demonstrate that protein workflows should not consider evolutionary information as a fixed characteristic of an alignment; evolutionary information can be tuned to shape downstream results.

## Methods

### Protein cohort, homologue retrieval, and parental MSAs

Our de novo generation analysis approach utilized 45 selected cohort proteins (Supplementary Data 1) with diverse taxonomies, functions, and sequence lengths. We retrieved the wild-type sequences from UniProt and refer to these as reference or anchor sequences hereafter. Each reference sequence was used for homologue retrieval from a local NCBI nr database downloaded on 16 August 2025.^19^ We retrieved homologues of reference sequences using NCBI BLASTP v2.13.0+^20^ where each query used the following parameters: maximum retrieved sequences set to 5,000, one high-scoring segment pair per target, and an E-value threshold of 1 × 10^-2. We combined the retrieved homologues with the original reference sequence followed by sequence deduplication. Sequences containing non-canonical amino acids were also removed. Using the remaining sequences, we generated parental MSAs for each cohort protein using MUSCLE v5.1 with Super5 mode.^21^

### Information targeting and matched depth control creation for de novo generative pipeline

Parental MSAs were organized so that the reference sequence was always pinned to index 0 in the parental alignment. We projected parental MSAs into the reference sequence coordinate frame such that only columns where the reference sequence contained a non-gap residue were used in the information-targeted and matched-depth control MSAs. Any gaps or non-canonical residues were encoded as zeros while canonical residues were encoded with tokens from 1-20.

We computed the per-column MSA information content using ProDy^22^ Shannon entropy. Because ProDy reports Shannon entropy using natural logarithms, bits/site was calculated as bits/site_j = log2(20) − H_j/ln(2), where H_j represents the Shannon entropy in nats (H) at a given column (j). We targeted MSA lower-tail information targets using the tenth percentile (p10) of per-column information values in the reference coordinate frame columns where at least 50% of the entries were non-gap tokens (50% occupancy also computed using ProDy). The information tuning procedure used a target grid of p10 values computed as log₂(20) − H_j, with H_j in nats, which we refer to as the pipeline coordinate. The grid spanned 1.90 to 3.10 in 0.05-unit increments in this coordinate. We computed the values in bits/site by converting to bits (via x/ln(2) − 1.913). The resulting search grid corresponds to approximately 0.83 to 2.56 bits/site. The operational threshold, 2.0 in the pipeline coordinate, was empirically chosen from the 45-protein cohort and corresponds to 0.972 bits/site.

The MSA-SNR row-coherence metric was computed as the mean of non-gap pairwise percent identity (PID) divided by the standard deviation of PID across tokenized sequences in the reference-projected coordinate frame. Cases where the metric became numerically unstable were not included in our MSA-SNR analysis.

The tuning process utilized a greedy-search procedure where non-reference rows were iteratively dropped to maximally increase or decrease MSA-SNR. An initial search direction was selected based on the starting p10 value and the desired p10 target. This search proceeded until 75% of the rows were dropped. If a p10 target was not achieved in one direction, the search was repeated in the opposite direction. A snapshot of the MSA’s p10 value in the pipeline coordinate was computed and stored after each row drop. If the process achieved the requested p10 target within a tolerance of 0.01 in the pipeline coordinate, then the row composition that provided the closest p10 value to the requested target was selected. In cases where a p10 target could not be achieved without dropping more than 75% of the rows in the MSA, that p10 target was skipped for that cohort protein. Due to the requirement that at least 25% of parental MSA rows must be maintained, only 20 cohort proteins produced MSAs on both sides of the 0.972-bit/site minimum threshold.

After creation of an information-targeted MSA, a matched-depth control MSA was created such that the reference protein and randomly selected rows were compiled to generate a control MSA with the same depth as the information-targeted MSA. As with information-targeted MSAs, the reference protein was pinned to index 0 and all protein sequences were tokenized and fixed inside the reference protein-projected coordinate frame.

Meff-style row weights were computed from the no-gap PID buffer with θ set to 0.70 for both information-targeted and control MSAs. We used these weights to create the downstream MSA context array.

### MSA context features and masked-diffusion model training

MSA context arrays were generated for each column to include:

- Meff-weighted amino acid frequencies
- gap frequency
- Shannon entropy
- non-gap residue occupancy
- inverse-sum-of-squares
- reference-residue probability
- one minus reference-residue probability
- indicator if reference and MSA mode match at each position
- MSA mode probability minus reference-residue probability

For each information-targeted or control MSA created, a separate MDLM denoiser^16^ was trained on the tokenized MSAs with the following vocabulary: gap tokens, 20 canonical amino acid tokens, mask token, and end of sequence token.

The study’s MDLMs represent tokenized sequences as learned positional and token embeddings and use the scalar corruption rate as a conditioning vector across all sequence positions. MDLM architecture used transformer blocks consisting of LayerNorm, followed by self-attention, optional cross-attention, and then feed-forward layers and dropout. MDLM parameters utilized a width of 256, a transformer block depth of 6, eight attention heads and feed-forward width of 1024, and a dropout rate of 0.1. We applied MSA-context cross-attention in the first and fourth transformer layers. For each step in the training process, non-gap positions were corrupted with corruption rates sampled from 0.50 to 0.95. Model training utilized an Adam optimizer (learning rate of 3 × 10^-4) where sparse categorical cross-entropy was computed over the masked positions for training and validation loss.

5% of the randomly shuffled MSA rows were selected as validation rows. Training was performed on the remaining rows using a batch size of 16. To mitigate the potential impact of MSA depth on MDLM training, we provided each model with a maximum training budget of 4,000 steps regardless of MSA depth. Validation scoring was performed every 200 steps across the validation rows using three seeded masks. After a burn-in period of 1,000 steps, a patience of 5 evaluation points (requiring an improvement of 1 × 10^-3) permitted models to stop before utilizing the full maximum training budget.

While training, classifier-free dropout with a probability of 0.1 replaced MSA context arrays with an array of zeros.

### De novo sequence decoding and candidate selection

We used a predefined grid of corruption rates for de novo sequence generation: 0.5, 0.65, 0.75, 0.85, 0.9 and 0.95. For each corruption rate, the reference sequence was corrupted, and 15 candidate sequences were generated across 12 decode steps. Each candidate sequence was generated using a deterministic seed. A cosine commit schedule was applied to a MaskGIT-style parallel decoding procedure.^23^ Gap, mask, and EOS tokens were forbidden during decoding to ensure that generated sequences would always maintain the reference projected coordinate frame.

Five sequences from each corruption rate were selected for downstream folding. However, we used latent representations from all 15 sequences for the latent geometry analysis shown in Fig. 4. We ranked each of the 15 sequences generated at each corruption rate according to their maximum percent identity to the full BLAST-derived sequences used for constructing the parental MSA. The generated-sequence selection approach utilized a novelty-forward procedure where the five sequences generated at each corruption rate with the lowest PID to the BLAST-derived sequences used for MSA construction were selected for downstream folding analysis. If PID ties occurred, such ties were resolved by utilizing a greedy k-center procedure in latent cosine space to select the most diverse five sequences.

### Structural analysis of generated de novo sequences

Chai-1 structure prediction^1^ of generated de novo sequences was always performed in an MSA-free manner. This design anticipated potential experimental confounds where an information-tuned MSA could steer not just de novo protein generation, but also the predicted structural outcomes. Structure prediction for each generated sequence was performed with ESM embeddings using the following parameter settings: 3 trunk recycles, 200 diffusion timesteps, and five generated protein models per input sequence.

Structural readouts were computed for each generated sequence using the per-Chai-1 model confidence outputs pLDDT, pTM, and PAE. We also folded each reference sequence with Chai-1 and used the TM-score binary^24^ to calculate the TM-score and RMSD between the predicted wild type and the generated sequence structures.

### MSA-SNR analysis and regime identification

Within Fig. 2, we performed analysis at the MSA cohort/model level to assess how trends in per-MSA information impacted generated-sequence novelty and structural confidence. Chai-1 maximum pLDDT was computed as the maximum of the mean pLDDT for each of the 30 sequences that were folded from each MSA. TM-score maximum was computed as the maximum TM-score achieved when comparing across Chai-1 protein models from each MSA’s 30 de novo sequences and the Chai-1 predicted structures for the MSA reference protein. Novelty was computed as 100% minus the maximum percent identity each generated sequence had to each of the BLAST-derived sequences that were used to construct the corresponding parental MSA.

We computed multiple information metrics after MSA generation, but p10 was the only Shannon information statistic that was used for information-targeting. From each created MSA, we computed the minimum bits/site content across all of the reference protein-projected columns. As in the tuning procedure, we determined this from ProDy-computed Shannon entropy after converting the entropy from nats to bits. However, in this case, the bits/site value was computed across all columns, not just those with an occupancy of at least 50%.

For each cohort protein, we identified the minimum bits/site value associated with the maximum MSA-SNR across the MSAs generated for that protein. Unweighted protein-cohort level MSA-SNR produced a mean peak location of 0.789 ± a standard deviation of 0.366 bits/site. Because the tuning procedure required at least a 25% row retention, we expected that the MSA-SNR maxima locations for each protein might not always reflect the true optimum. To give more weight to MSA-SNR maxima that were higher, we computed a Softmax-weighted peak of 0.964 ± 0.123 bits/site (temperature of 1.0). Thus, this reference point was empirically calculated from the 45-protein cohort dataset and was derived independently from the p10 tuning process. In the paper, when we refer to ‘below-threshold’ MSAs or downstream artifacts, this means that the generated MSA had a minimum information content of < 0.972 bits/site. ‘Above-threshold’ indicates that the generated MSA had a minimum information content of ≥ 0.972 bits/site. We generally describe this operational threshold as approximately 1 bit/site.

In the confidence-novelty analyses, we computed the mean Chai-1 pLDDT for sequences generated from each MSA or the mean TM-score from each structure generated from each MSA. For each generated sequence, we also computed a novelty fraction value, which we define as 1 minus the maximum PID ratio of the BLAST-derived sequences used to construct the parental MSA. The structural confidences were multiplied by the novelty fraction values for each of the 30 sequences that were folded from each MSA. The reported maximum novelty-confidence values were then the maximum of the structural confidence metric multiplied by the novelty fraction value for the 30 sequences that were folded from each MSA.

MSA-SNR sensitivity analysis assessed how the mean PID numerator and PID dispersion denominator contributed to the peak located near the 1-bit/site operational threshold, as well as how these two components correlated with the structural, novelty, and novelty-frontier metrics. To keep the denominator comparison the same sign as in the full metric, PID dispersion was converted to negative PID dispersion. We also performed partial Spearman analyses with Meff (θ=0.70) and MSA depth to determine how much of the relationship between MSA-SNR and the folding/novelty/frontier metrics could be explained by Meff and depth (Supplementary Data 6).

We speculated that the correlations between MSA-SNR and confidence/novelty outcomes might be fragile or driven by a small subset of the cohort proteins. To test this, we performed 1,000 deterministic replicates where only 36 of the 45 proteins were sampled to compute the Spearman correlations. This gave us an estimate of how stable the correlations were (Supplementary Data 6) as protein dataset composition was perturbed.

### MSA analysis

In Fig. 3 and 4 plots, we display only values from the information-tuned alignments as each of the matched control MSAs inherited a below- or above-threshold label from a matched tuned MSA that did not necessarily reflect the bits/site minimum of the control MSA. ProDy was used to calculate the MI-APC top-quantile mean metric (the mean APC-corrected mutual information for values in the top quantile).^25^

We constructed MSA-similarity graphs such that edges connected any rows where the pair’s no-gap PID was ≥ 0.70. NetworkX^26^ was used for computing largest connected component (LCC) fraction, edge density, average shortest path length within the LCC, bridge and articulation fractions. We used the NetworkX Laplacian matrix to compute graph connectivity *λ*_2_ on the LCC from the sequence similarity graph using SciPy^27^ sparse eigsh where we report the second-smallest eigenvalue.

The parameterized schematic in Fig. 3a provides a visual representation of protein-level median values below and above the operational minimum-information threshold of 0.972 bits/site. This is not meant to represent an actual graph from the dataset, but rather to provide an intuitive visual representation of dataset trends. For the below- and above-threshold schematics, we generated synthetic graphs by assigning a fixed number of nodes to each graph where the number of clusters could vary. Relative cluster structure was determined by the protein median component count and LCC fraction while edge density and *λ*_2_ set the number of edges that occur inside and between clusters. The LCC path length determined the size of the graph while the fragility metrics (bridge and articulation fractions) were used to set the number of bottlenecks in the graph.

### Latent analysis

For Fig. 4 analysis, we calculated the distance between each of the generated-protein latent representations and the anchor latent representation (from the reference protein) in the original latent space as Euclidean distance. All other downstream metrics were computed following mean-centering of the latent representations.

We measured local kNN radius using the scikit-learn NearestNeighbors where the metric shown in Fig. 4 represents the median distance across k=5. Latent graph representations were generated using scikit-learn kneighbors_graph with k=15. After symmetrizing each graph, we used the NetworkX library to compute clustering and the Gini coefficient. Graph connectivity *λ*_2_ was computed using the same methodology as in Fig. 3.

We calculated hubness directly from the directed kNN neighbor lists by quantifying how often each latent representation occurred in the neighbor list of each other latent representation. The hubness metric we report is the standard deviation of these values.

The scikit-learn implementation of Ledoit-Wolf shrinkage covariance^28^ was used to compute covariance metrics. The computed covariance matrices were symmetrized before calculating participation ratio (PR) from the covariance eigenvalues. Sphericity was computed as PR divided by the latent space dimensionality. Spectral entropy was computed from the eigenvalues, and effective rank was computed from the exponential of the spectral entropy.

As in Fig. 3a, Fig. 4a provides a parameterized schematic of cohort-protein-level median values on either side of the operational minimum-information threshold of 0.972 bits/site. The representations are generated from two-dimensional point clouds. The sizes of the point clouds were set by the size of the local kNN radii. The sphericity metric controls the anisotropy of the cloud while *λ*_2_ and clustering control edge density. The Gini coefficient and hubness set hub points. The anchor latent representation (reference protein) is represented as a black dot in the center of each point cloud.

For the metrics computed and compared in Figs. 3 and 4, we also performed a paired analysis for the 20 cohort proteins that produced MSAs on both sides of the operational threshold of 0.972 bits/site. We compared the protein-level mean values on either side of the threshold using a two-sided paired t-test and corrected p-values with a Benjamini-Hochberg correction. We also replicated the inter-feature Spearman analyses shown in Fig. 3b and 4b using only the 20 proteins that were represented on both sides of the operational threshold (Supplementary Data 6).

Violin plot comparisons in Fig. 3 and 4 were made using two-sided Welch tests and corrected using Benjamini-Hochberg q-values. We computed the BH correction for each individual metric using Welch tests between the information-targeted below- and above-threshold protein means and between control MSA below- and above-threshold protein means. This approach usually provided small statistical corrections but allowed each correction to be metric focused. In Supplementary Data 6, we also show the same statistical analyses at minimum-information thresholds of 0.8, 0.9, 0.972, 1.0, 1.1, and 1.2 bits/site.

### Model dynamics analysis

To examine per-residue model dynamics, we compiled per-residue data for each of the de novo generated sequences that were selected for folding. We determined if a residue was mutated based on whether it differed from the reference sequence at the corresponding position. We measured the MSA bits/site composition at the corresponding positions (using ProDy as previously described) for each residue in generated sequences as well as the Chai-1 pLDDT for predicted structures. We determined negative log-likelihood (NLL) with and without conditioning by providing each MDLM with the corrupted reference sequence states with and without the MSA context array to determine the scoring-time utility of the context array. ΔNLL was defined as unconditioned NLL minus conditioned NLL such that positive values indicate that the MSA context improved model confidence. Decoder conditional entropy was computed from MDLM predictive distributions when MSA context was present and is distinct from MSA Shannon entropy.

Even though these analyses occurred at the residue level, we aggregated the residue-level data as with protein-level medians to prevent longer sequences or cohort proteins that provided more de novo sequences from being overweighted in the results.

The mutation analysis in Fig. 5a assigns mutation rates to MSA bits/site bins and computes mutation probability from the protein-level summaries within that bin. The decode-step analyses in Fig. 5b-e use the median of protein-level means for decode step (Fig. 5b-d) or cumulative medians of protein-level means at successive decode steps (Fig. 5e). Fig. 5f compares matched target-minus-control outcomes where results are first averaged at the protein level such that the violin plots are showing protein-level paired differences.

In Supplementary Fig. 3, we analyzed post hoc conditioned and unconditioned confidence across MSA-confidence bins using the same MDLM models and corrupted states used in sequence generation. Jensen-Shannon divergence was computed between the per-residue conditioned/unconditioned distributions. Supplementary Fig. 3c,d examine conditioned uncertainty scores computed from decoder entropy and NLL. These results compared how well model uncertainty was predictive of whether Chai-1 pLDDT was at/above or below 70. The Brier-style analysis shows rank-normalized scores within each protein and condition before comparing to the Chai-1 pLDDT outcome. Aggregation of values in the foldability-surface decile bins represents protein-level means for each decile of mutation rate or decoder uncertainty.

### Information flow analysis

The examples shown in Fig. 6a were selected based on the following criteria: We selected from MSAs where the reference protein length was at least 100 residues and all sequences generated from the MSA had at least a downstream mean Chai-1 pLDDT of 70. From this pool of MSAs, we selected the within-protein pair of MSAs with the greatest difference in average bits/site information content, requiring that one MSA was below the operational minimum-information threshold of 0.972 bits/site and the other was at or above the threshold. We then selected each MSA’s sequence with the greatest novelty relative to the BLAST-derived sequences used to construct the parental MSA. We displayed raw residue traces with an overlaid rolling mean trend using a nine-residue window size. Residues that were mutated relative to the reference sequence are noted by rug marks on the plot.

For the local information flow analysis shown in Fig. 6b, we computed per-residue Spearman correlations within each of the sequences selected for folding. As in Fig. 5, we aggregated the data as protein-level medians to prevent uneven representation by proteins with longer sequences or higher generated sequence counts. The residue-level confidence band plots showing pLDDT and mean PAE probabilities in Fig. 6c,d and Supplementary Fig. 4b,c represent aggregated protein-level probabilities within each MSA bits/site bin. Fig. 6c,d represent all residues while Supplementary Fig. 4b,c represent only residues where a mutation occurred.

### Mediation-style pathway screening analysis

We performed a mediation-style pathway screen to assess how information correlates above and below the operational minimum-information threshold of 0.972 bits/site. We do not infer causal mediation through specific pathways; this analysis simply demonstrates the presence of reproducible covariance chains within the selected metrics. The mediation screen occurred at the MSA-level starting from the median bits/site content of each MSA. We then screened for associations in the Fig. 3 feature set, then the Fig. 4 feature set, and finally to median Chai-1 pLDDT or PAE values as the end points in the chain. We separated information-targeted MSAs by orientation to the threshold and then assessed covariance over MSA-level medians.

For above- and below-threshold information-targeted MSAs, we computed a serial indirect-effect score from standardized linear models moving from MSA information to MSA mediator to latent mediator to Chai-1 pLDDT or PAE. Analysis across these points produced three coefficients. The serial indirect-effect score was their product. We computed confidence intervals using 5,000 bootstrap resamples.

We considered that covariance might be dominated by protein-level differences, so we performed protein-level grouped cross-validation and required that at least four of five validation folds have the same sign with indirect effect scores more than 0.005. In addition, we required that successful pathways have bootstrap confidence intervals that exclude zero.

### Independent structure-prediction benchmark analysis

We initially constructed an independent structure dataset where all structures were deposited in the RCSB database after 01 January 2023. These structures were required to have a resolution ≤2.0 Å, no more than one polymer entity, only one biological assembly, and no more than 20% of the residues could be missing from the deposited structure. Only canonical amino acids were permitted, except for selenomethionine which we converted to methionine. Diversity was enforced by keeping only one monomer and one multimer (of a single polymer) from each 30% sequence-identity cluster ID in the RCSB database API.

Our benchmark JackHMMER^17^ MSAs were generated from a local UniRef90 database accessed 14 Jun 2022. After running JackHMMER for three iterations, the final output was converted to A3M format. The MSV, Viterbi, and Forward filter thresholds were 5 × 10^-4, 5 × 10^-5, and 5 × 10^-7, respectively. Reporting and inclusion E-value thresholds were both 1 × 10^-4.

Before tuning JackHMMER alignments to specific fractions of columns below the operational threshold of 0.972 bits/site, we removed dot characters and kept only reference-protein-projected columns where the query sequence was a non-gap value. We also dropped rows with > 25% gaps and deduplicated using SimHash buckets. Because constructing the PID buffer becomes very memory intensive with deep alignments, we downsampled to 20,000 rows in cases where the depth was higher than 20,000 sequences.

As when we tuned MSAs for protein generation, we performed a bi-directional MSA-SNR sweep. Here, we did not impose a maximum percentage of rows that could be removed. The approach greedily increased and decreased MSA-SNR as much as possible by removing rows starting from the processed JackHMMER MSAs. Column-wise bits/site, after conversion of ProDy entropy from nats to bits, and Meff were computed after each row removal. When possible, the MSA that most closely achieved the target fraction of columns below the operational threshold of 0.972 bits/site within a tolerance of 1 percentage point was retained for downstream analysis. In the case of ties, we chose the alignment with the most sequences. The fraction targets shown in Fig. 7 are 0, 2, 5, 7.5, 10, 12.5, 15, 20, 25, and 30%. We attempted higher fractions not shown in Fig. 7 but found that the number of MSAs that could achieve fractions higher than 30% dropped too much for robust analysis.

Chai-1 structure prediction was performed with either the JackHMMER or tuned MSA provided as a local input. No external templates were used. TM-score and RMSD were computed using the TM-score binary. Our initial dataset was composed of 1,178 experimental structures but was reduced to the 441 structures that could produce MSAs across all targeted fractions and for which we did not observe TM-score errors or missing values.

In Fig. 7, we defined deltas such that a positive value would indicate that the tuned MSA displayed a more accurate structure prediction than the original JackHMMER MSA. Mean deltas were computed within each of the ten targeted fractions, with 95% confidence intervals computed using 10,000 bootstrapping iterations. ΔpTM-gated analysis retained only comparisons where the Chai-1 pTM was higher for the tuned MSA by the indicated threshold. We chose ΔpTM > 0.03 as a primary gate as it represented a moderate confidence increase where there were still enough structures for analysis across each target fraction. In the Fig. 7g visualization, we superimposed Chai-1 predicted structures onto the experimental structure using PyMOL. MSA columns with bits/site <0.972 were colored pink, while columns with bits/site ≥0.972 were colored blue. The experimental structure was rendered in partially transparent grey.

In Supplementary Fig. 5c,d, we display local-error transition analysis where we compare structural accuracy for paired JackHMMER-derived and targeted-fraction MSAs against the experimental structure. Each Chai-1 inference produced five predicted structures. We selected whichever had the highest pTM score for this analysis. For each selected Chai-1 structure, we aligned the predicted Cα residues with the experimental Cα residues using Kabsch^29^ alignment. We considered local information and error based on windows where each was composed of a central residue and the five residues on either side. For each generated window, we assessed whether the mean bits/site content of the window was below or at/above the operational threshold of 0.972 bits/site in the JackHMMER and tuned alignments. We assigned four classes to our windows to indicate how each window changed from the JackHMMER MSA to the tuned MSA: above→above, above→below, below→below, and below→above. We then stratified each of these transition states into structure error bands according to whether the local error improved or worsened due to tuning. We averaged across target fractions for each PDB and balanced PDB representation so that PDBs contributed as equally as possible. In Supplementary Fig. 5d, we applied the gate, ΔpTM > 0.03, to illustrate the amplification of local error improvements that occurred when the tuning process improved Chai-1 confidence.

### Statistical units and reproducibility

Fig. 2 analyses report statistics over MSA-level data, while Figs. 3 and 4 compare the mean representations for each cohort protein to avoid treating individual MSAs or sequences as independent biological replicates. Figs. 5 and 6 both analyze residue-level trends, but they aggregate these results at the cohort-protein level to give each cohort protein equal weight.

Figures were generated using Matplotlib^30^ and seaborn^31^ Python libraries. The source files and metadata are provided in the project repository.

Across these experiments, deterministic seeds were used in all stochastic processes such as row-shuffling, generation of random-row control MSAs, model training, de novo sequence generation, and bootstrapping to preserve reproducibility.

Scripts invoking Chai-1 were run in a Chai-1-specific environment containing chai_lab v0.6.1, TensorFlow v2.19.0, PyTorch v2.6.0, NumPy v1.26.4, SciPy v1.15.3, pandas v2.2.3, scikit-learn v1.6.1, NetworkX v3.4.2, Biopython v1.85, Biotite v1.4.0, Gemmi v0.6.7, Matplotlib v3.10.3, PyArrow v20.0.0, and OpenPyXL v3.1.5.

Non-Chai-1 scripts were run in a separate analysis environment containing TensorFlow v2.8.0, PyTorch v2.8.0, NumPy v1.23.5, SciPy v1.10.1, pandas v2.3.2, scikit-learn v1.2.1, ProDy v2.4.1, scikit-dimension v0.3.4, factor-analyzer v0.5.1, NetworkX v2.5.1, Biopython v1.79, seaborn v0.13.2, Matplotlib v3.9.4, PyArrow v10.0.1, and OpenPyXL v3.1.5.

## Data availability

The scripts and data tables necessary to reproduce the manuscript results and figures are deposited at github.com/Evotec-in-silico-AI/MSA-information-tuning. The data files needed to reproduce this study and replicate the described analysis are available from Zenodo repository 10.5281/zenodo.23671811.

## Code availability

The codebase used to generate the results described in this manuscript is deposited at github.com/Evotec-in-silico-AI/MSA-information-tuning along with sample data and instructions for reproducing the experiments and downstream analyses.

## Acknowledgements

G.R. acknowledges Evotec for institutional support and computational resources. The authors wish to thank Chandrajit Bajaj for his excellent advice.

## Author contributions

G.R. conceived the study, developed the analysis framework, performed MSA tuning, model training, structural analyses, figure generation and manuscript drafting. L.C. provided scientific supervision and contributed to manuscript review.

---

**Supplementary Fig. 1.**
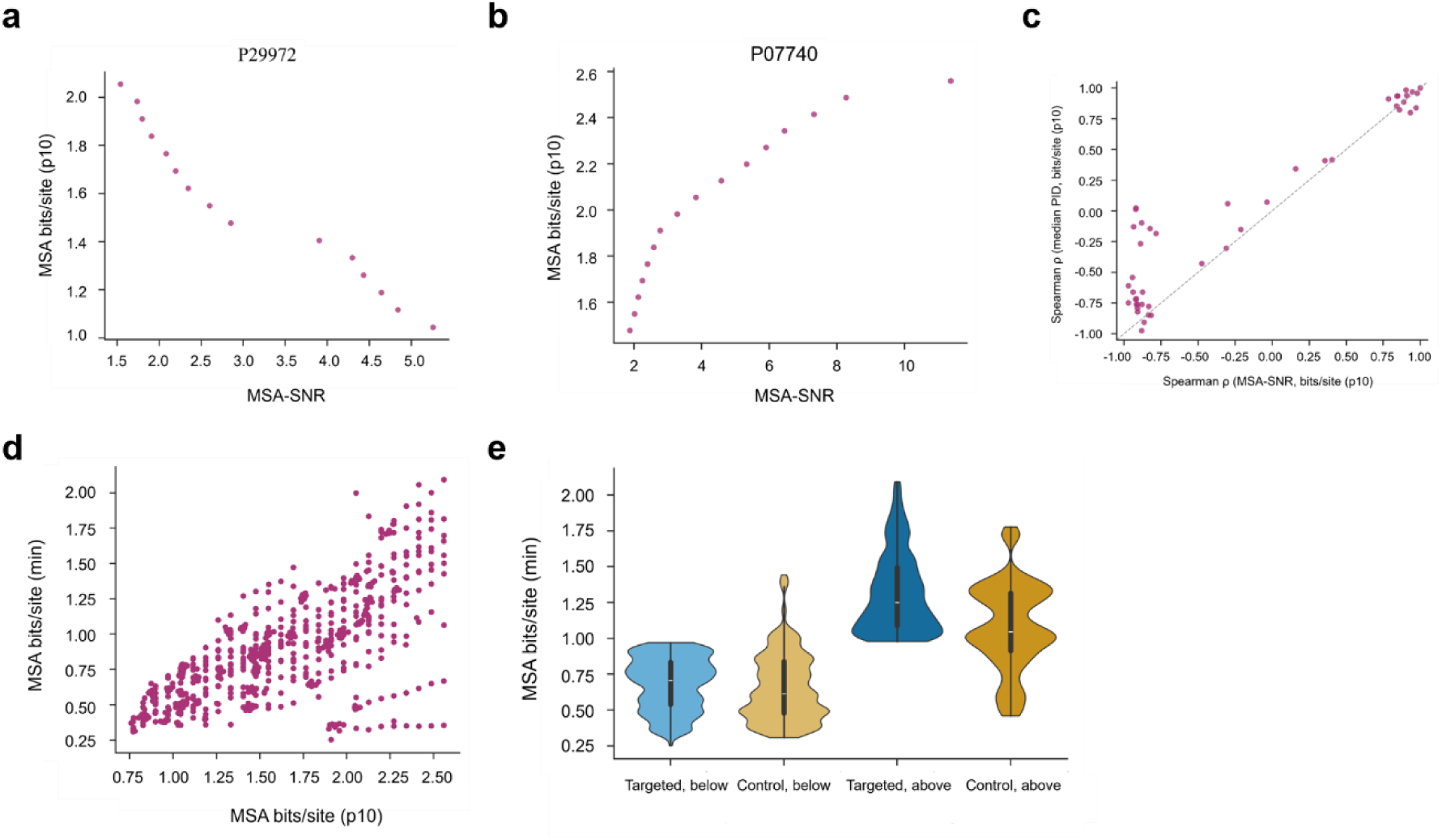
MSA-SNR varies non-monotonically with lower-tail MSA information. MSA-SNR is shown relative to p10 and minimum bits/site. p10 is the lower-tail information metric used for information tuning, while MSA minimum bits/site is the metric used as a threshold throughout the manuscript. **a**, Example targeted-MSA trajectory for P29972, where MSA-SNR decreases as p10 increases. **b**, Example targeted-MSA trajectory for P07740, where MSA-SNR increases as p10 increases. **c**, Protein-level comparison of Spearman correlations between MSA-SNR and p10 versus median pairwise identity and p10. Proteins in which median pairwise identity increases with p10 tend to show a more positive MSA-SNR-p10 relationship. **d**, Relationship between p10 and minimum bits/site across MSA samples. These lower-tail information summaries are positively associated but not identical. **e**, Distribution of minimum bits/site across downstream condition groups. Labels are assigned from information-targeted MSAs relative to whether their minimum bits/site value was above or below threshold. Matched controls inherit the label of their paired targeted MSA. For this reason, their minimum bits/site value can fall on either side of the threshold regardless of label assignment.

**Supplementary Fig. 2.**
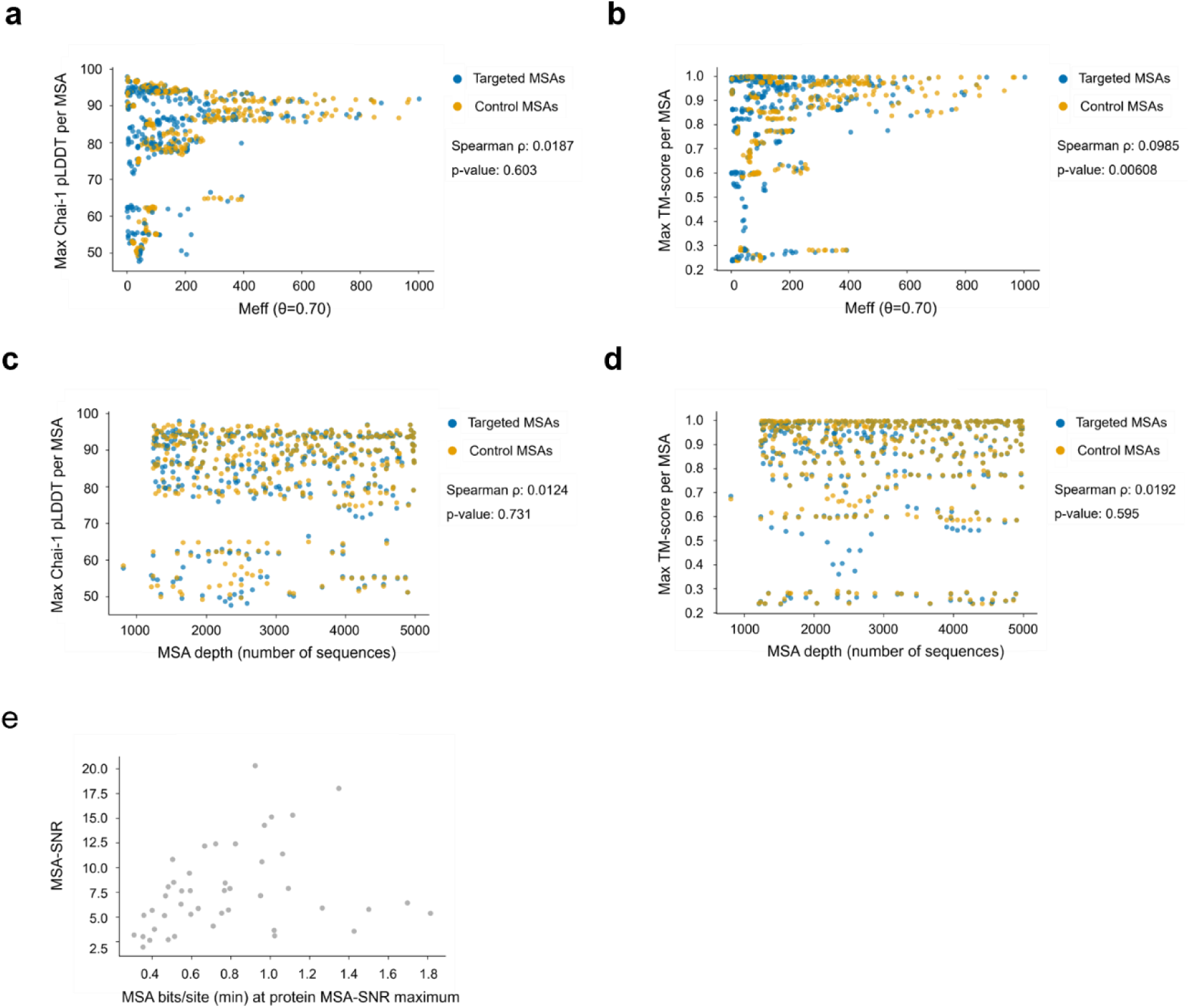
Meff and retained depth do not reproduce the MSA-SNR regime. **a**, Maximum mean Chai-1 pLDDT across generated sequences from each MSA plotted against Meff (Spearman ρ = 0.0187, p = 0.603, n = 774). **b**, Maximum mean TM-score to the Chai-1-predicted anchor structure plotted against Meff (Spearman ρ = 0.0985, p = 0.00608, n = 774). **c**, Maximum mean Chai-1 pLDDT across generated sequences from each MSA plotted against retained MSA depth (Spearman ρ = 0.0124, p = 0.731, n = 774). **d**, Maximum mean TM-score to the Chai-1-predicted anchor structure plotted against retained MSA depth (Spearman ρ = 0.0192, p = 0.595, n = 774). **e**, Per-protein MSA-SNR maxima plotted against the minimum bits/site value at which that peak is observed. This panel isolates per-protein MSA-SNR maxima from the full MSA-level point cloud and complements the peak-location summaries in Fig. 2e**,f**.

**Supplementary Fig. 3.**
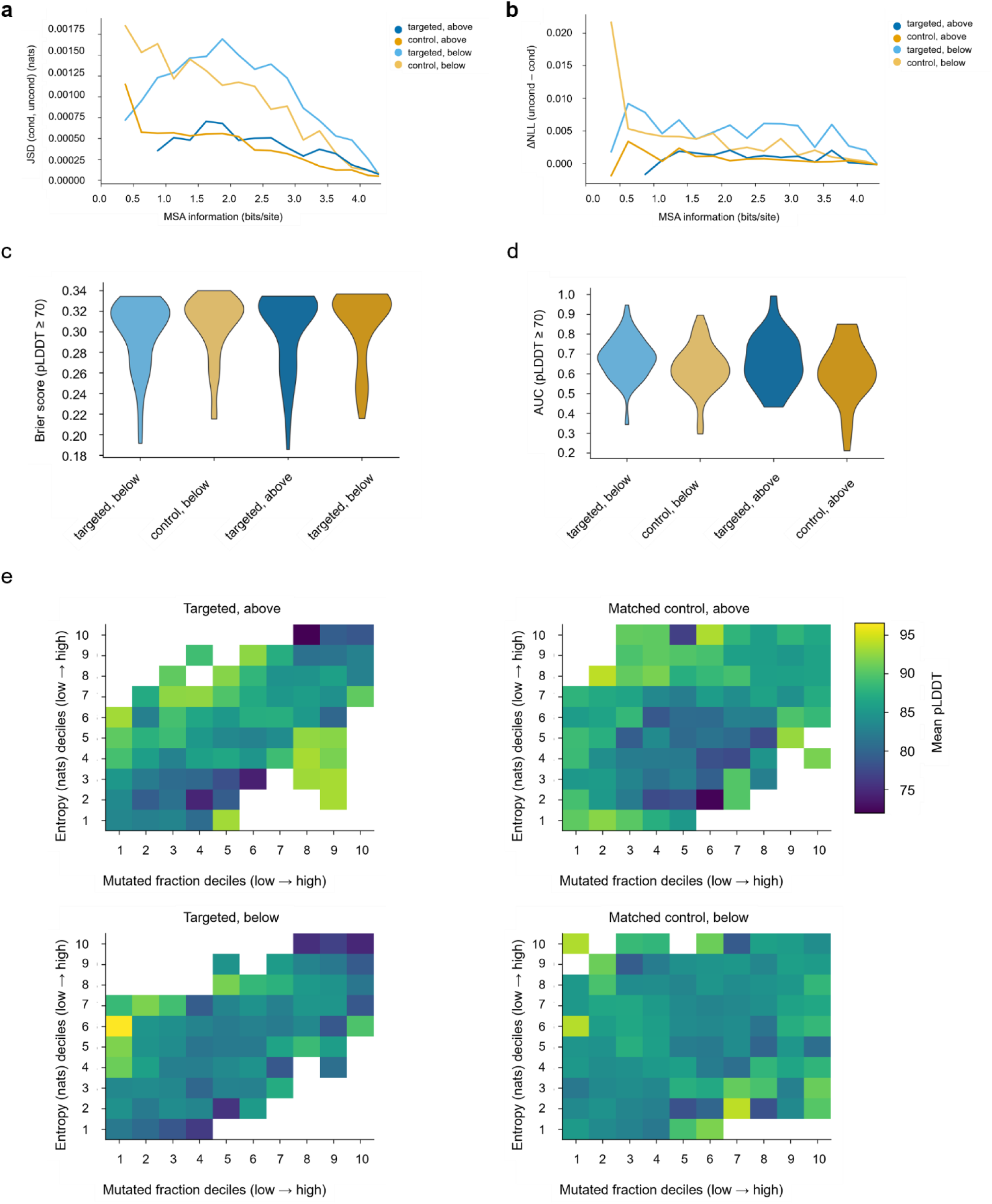
MSA context sensitivity is strongest at low-information positions. **a**, Jensen-Shannon divergence (JSD) between per-site predictive distributions evaluated with and without MSA context, plotted as a function of local MSA information. Higher JSD indicates a larger MSA context-dependent change in the MDLM predictive distribution. **b**, Change in per-site NLL between scoring without and with MSA context. ΔNLL is defined as NLL_uncond − NLL_cond, so positive values indicate improved likelihood for the generated residue when MSA context is supplied. **c**, Brier-style score for predicting Chai-1 pLDDT ≥ 70 from conditioned uncertainty scores. Lower values indicate stronger agreement between uncertainty ranking and structural-confidence outcome. **d**, Protein-level area under the receiver operating characteristic curve (AUC) for the same pLDDT ≥ 70 prediction task using conditioned uncertainty scores. Higher AUC indicates better discrimination between residues above and below the pLDDT threshold. **e**, Structural-confidence surfaces across mutation fraction deciles and MSA-context conditioned decoder entropy deciles. Grid cells are aggregated at the cohort protein-mean level.

**Supplementary Fig. 4.**
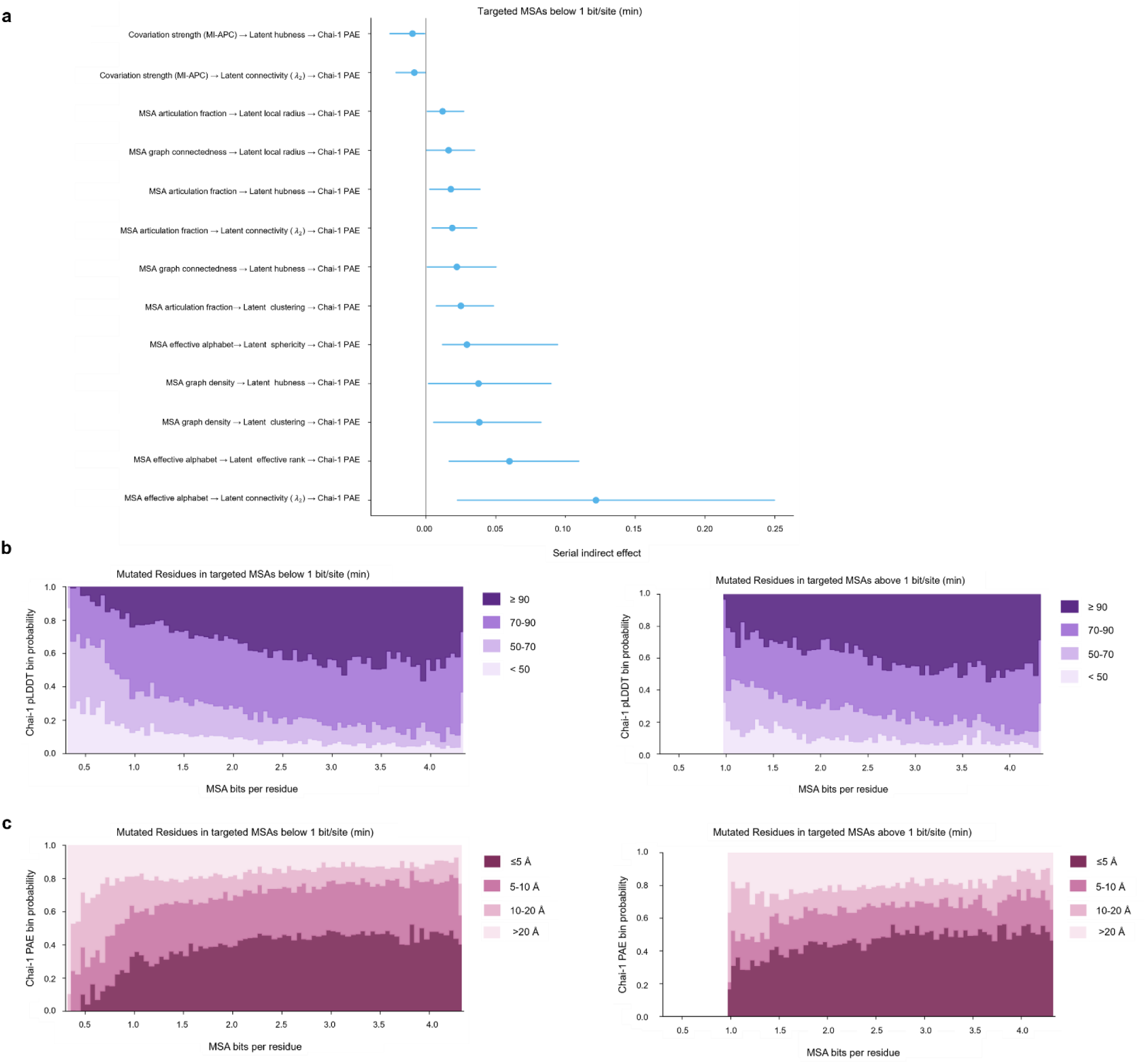
Lower-information MSAs show enriched MSA-to-latent-to-PAE covariance structure and reduced confidence at mutated residues. **a**, Thirteen putative covariance chains survive in the lower-information regime. The strongest lower-information chain is MSA effective alphabet → latent connectivity λ₂ → Chai-1 mean PAE, with standardized serial indirect effect 0.1218 and 95% bootstrap confidence interval 0.0225–0.2497. No chains survive in the higher-information regime. **b**, Protein-balanced Chai-1 pLDDT confidence distributions for mutated residues across local MSA bits/site bins in lower- and higher-information targeted samples. Residues are assigned to fixed-width local bits/site bins of 0.05 and summarized over pLDDT bands <50, 50-70, 70-90, and ≥90. **c**, Protein-balanced per-residue mean PAE distributions for mutated residues across the same local MSA bits/site bins. Residues are summarized over PAE bands ≤5, 5-10, 10-20, and >20.

**Supplementary Fig. 5.**
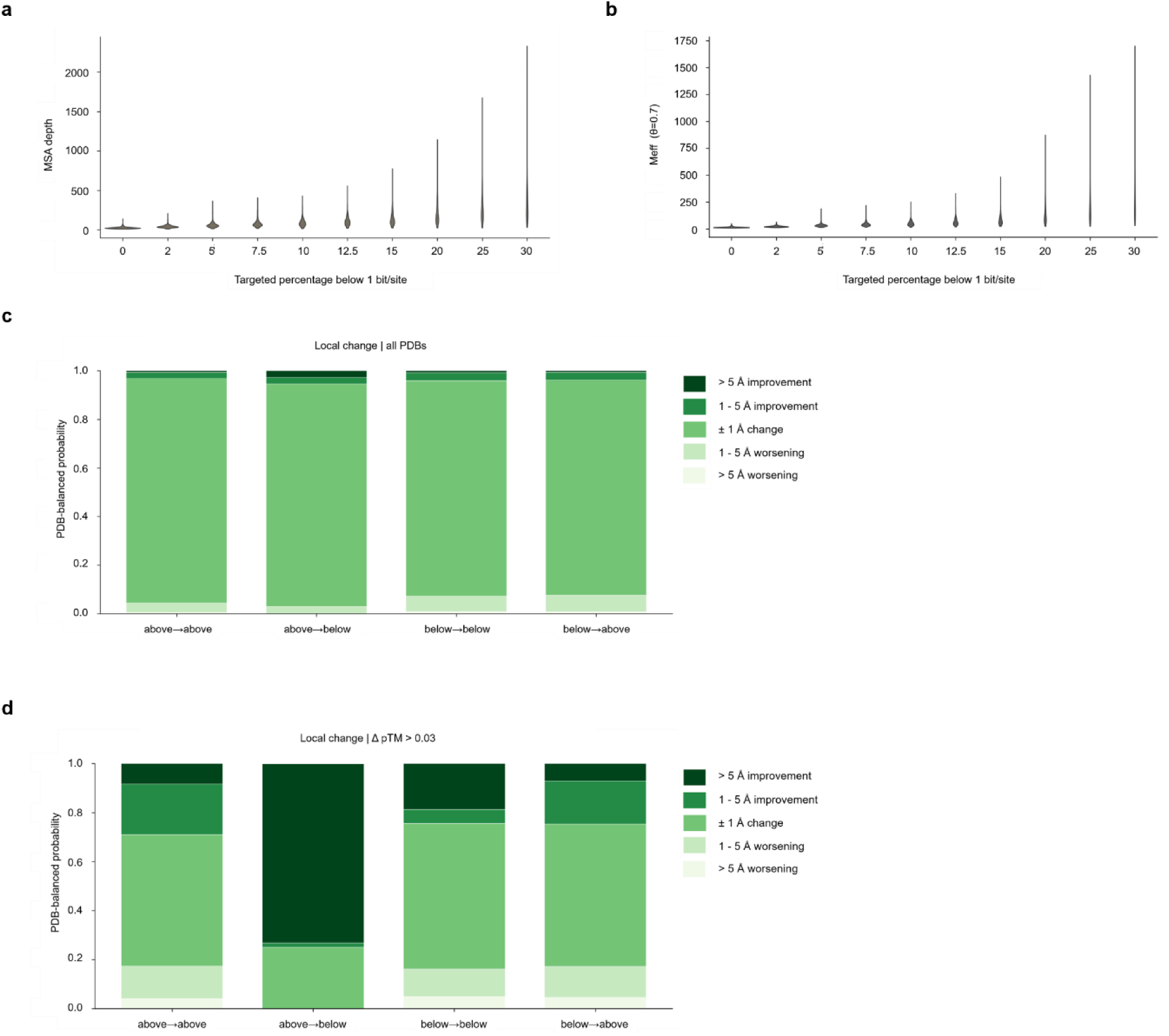
Alignment size increases with below-reference fraction but does not explain confidence-gated folding gains. **a**, Retained MSA depth across targeted fractions below the 1-bit/site threshold. **b**, Effective sequence count (Meff at θ = 0.70) across the same target fractions. **c**, Local Cα-error changes across information-transition classes without confidence gating. Residue-centered windows are classified by whether JackHMMER and tuned centered-window bits/site lie below or at or above the 0.972-bit/site threshold (above→above, above→below, below→below, below→above). Local structural outcomes are summarized as fractions of residue windows falling into improvement and worsening bands (>5 Å improvement, 1–5 Å improvement, ±1 Å change, 1–5 Å worsening, >5 Å worsening). **d**, Local Cα-error changes across the same transition classes after applying the ΔpTM > 0.03 confidence gate.

